# The Ran pathway uniquely regulates cytokinesis in cells with different fates in the early *C. elegans* embryo

**DOI:** 10.1101/2021.01.06.425598

**Authors:** Imge Ozugergin, Karina Mastronardi, Chris Law, Alisa Piekny

## Abstract

Cytokinesis occurs at the end of mitosis and occurs due to the ingression of a contractile ring that cleaves the daughter cells. This process is tightly controlled to prevent cell fate changes or aneuploidy, and the core machinery is highly conserved among metazoans. Multiple mechanisms regulate cytokinesis, but their requirement in different cell types is not known. Here, we show that differently fated AB and P_1_ cells in the early *C. elegans* embryo have unique cytokinesis kinetics supported by distinct levels and cortical patterning of myosin. Through perturbation of polarity regulators and the generation of stable tetraploid strains, we demonstrate that these differences depend on both cell fate and size. Additionally, these parameters could influence the Ran pathway, which coordinates the contractile ring with chromatin position, and controls cytokinesis differently in AB and P_1_ cells. Our findings demonstrate the need to consider multiple parameters when modeling ring kinetics.

## INTRODUCTION

We have extensive knowledge of the core machinery that regulates cytokinesis, but the mechanisms that regulate this machinery are less well-understood. Cytokinesis occurs during mitotic exit due to the ingression of a RhoA-dependent contractile ring that assembles in the equatorial plane. The Rho guanine nucleotide exchange factor (GEF) Ect2 (*C. elegans* ECT-2) generates active RhoA in the equatorial plane, which in turn directs the assembly of a contractile ring by recruiting effectors for F-actin polymerization and myosin activation. Consistent with its essential role in this process, Ect2 depletion causes cytokinesis failure in multiple cell types and organisms (Chalamalasetty et al., 2006; Dechant & Glotzer, 2003; Echard & O’Farrell, 2003; Kotynkova et al., 2016; Nishimura & Yonemura, 2006; Prokopenko et al., 1999; Tatsumoto et al., 1999; Yuce et al., 2005; Zhao & Fang, 2005). In early anaphase, actomyosin filaments assemble as a broad equatorial band, then transition into a tight ring that pinches in the overlying cortex (Green et al., 2012; Lewellyn et al., 2010; van Oostende Triplet et al., 2014). Various proteins control ring closure kinetics via crosslinking actin and/or regulating myosin activity. The highly conserved protein anillin (*C. elegans* ANI-1) is a key regulator of cytokinesis, and anchors the contractile ring to the membrane through binding sites for RhoA, F-actin, myosin, microtubules and phospholipids (Piekny & Maddox, 2010; Tse et al., 2011; van Oostende Triplet et al., 2014). Numerous mechanisms regulate cytokinesis, but their relative contribution in different cell types is not well-understood, since few studies have been done in comparable cell types (*e*.*g*. cells with different fates in the same organism) (Green et al., 2012; Piekny et al., 2005; Pollard & O’Shaughnessy, 2019).

The concerted action of multiple pathways ensures robust and efficient cytokinesis. Signals from microtubule-independent locations of the cell such as the cortex, centrosomes, kinetochores and/or chromatin have been shown to regulate cytokinesis, although their conservation and relative contributions are not well-defined (Beaudet et al., 2017; Beaudet et al., 2020; Cabernard et al., 2010; Dechant & Glotzer, 2003; Deng et al., 2007; Kiyomitsu & Cheeseman, 2013; Petronczki et al., 2007; Rodrigues et al., 2015; Silverman-Gavrila et al., 2008; Zanin et al., 2013). These pathways may function redundantly with spindle-dependent pathways in symmetrically dividing cells, but can also be essential depending on factors such as cell size and fate.

Cues associated with chromatin coordinate the position of the contractile ring in relation to segregating chromosomes in HeLa cells, but it is not known if this mechanism functions in other organisms and cell types (Beaudet et al., 2017; Beaudet et al., 2020; Kiyomitsu & Cheeseman, 2013). The small GTPase Ran is activated by RCC1 (RanGEF; *C. elegans* RAN-3) near chromatin and is inactivated by RanGAP (*C. elegans* RAN-2) in the cytosol. Following nuclear envelope breakdown in mitosis, a gradient of Ran-GTP forms with high levels near chromatin, and low levels near the cortex (Clarke & Zhang, 2008; Kalab et al., 2006; Kalab et al., 2002). A heterodimer of importin-α/β binds to the nuclear localization signal (NLS) of proteins, which are released by Ran-GTP. Since importin-binding can influence protein function, NLS-containing proteins could be differently regulated at the cortex vs. in the vicinity of chromatin. In support of this, meiosis of mouse oocytes requires cortical polarization to form the polar body, which occurs at specific distances to chromatin and is controlled by Ran-GTP (Deng et al., 2007). During cytokinesis, importin-β binds to and is required for the localization and function of anillin in HeLa cells by enhancing its recruitment to the equatorial cortex during anaphase (Beaudet et al., 2017; Beaudet et al., 2020). The Ran-dependent regulation of cytokinesis has not been studied in other cell types and we propose that it could play an important role in furrow positioning in cells where chromatin is positioned closer to the cortex.

Cytokinesis has been well-characterized in the *C. elegans* P_0_ zygote, which is influenced by anterior-posterior polarity. This cell divides asymmetrically to give rise to a larger, anterior AB cell whose descendants form multiple tissues, and a smaller, posterior P_1_ cell fated to become the germline (Rose & Gonczy, 2014). Anterior-posterior polarity is controlled by the mutually exclusive distribution of anterior (PAR-3/PAR-6/PKC-3) and posterior (PAR-2/PAR-1) complexes along the cortex. The establishment of polarity depends on the asymmetric enrichment of actomyosin contractility, which occurs in response to sperm entry (Cowan & Hyman, 2007; Rose & Gonczy, 2014). Polarity is maintained via feedback between the PAR proteins and the actomyosin system at the anterior cortex, although its control switches from regulation by RhoA to Cdc42 (Cowan & Hyman, 2007). As the P_0_ zygote enters anaphase, actomyosin appears as patches or clusters at the anterior and equatorial cortex (Munro et al., 2004; Tse et al., 2012). Compression-driven flows at the equatorial cortex may help actomyosin filaments to accumulate and align to mediate asymmetric ring closure (Khaliullin et al., 2018; Maddox et al., 2007). Three phases of ring closure have been defined, although it is not clear how flows influence these phases: 1) ring assembly, with little to no ingression of the membrane, followed by 2) furrow initiation, when the membrane is indented by the contractile ring, and 3) constriction, when the membrane ingresses until it reaches the midbody [e.g. (Chan et al., 2019; Khaliullin et al., 2018; Lewellyn et al., 2010; Osorio et al., 2019; Price & Rose, 2017)]. Parameters such as size and fate influence ring closure kinetics. For example, there is a negative correlation between constriction and cell size, which was proposed to help coordinate the timing of cytokinesis among different cells during embryogenesis (Carvalho et al., 2009). Cell fate also may play an important role as Davies et al. (2018) showed that different cells in the 4-cell embryo exhibit unique kinetics compared to the P_0_ cell, attributed to extrinsic and intrinsic differences in regulating formin-derived F-actin.

In this study, we show that AB and P_1_ cells have unique ring closure kinetics that are governed by cell fate and size, and are mediated by different Ran pathway components. We found while AB cells have rapid ring assembly and initiation, these phases are slower in P_1_ cells. In support of this difference, AB cells have higher levels and greater alignment of myosin at the equatorial cortex, along with stronger cortical flows compared to P_1_ cells. In line with their cortical flows, we found that the ring closes asymmetrically in AB cells, but not in P_1_ cells. We show that these differences are due to cell fate and size, which reflect distinct threshold levels and organization of myosin. Disrupting cell fate via depletion of PAR-1 or PAR-3 equalizes kinetics, which are similar to AB cells. Increasing cell size, via the generation of tetraploid strains where AB and P_1_ cells are properly fated but larger in size, causes P_1_ cells to have ring closure kinetics and myosin distribution that is similar to diploid AB cells. Since AB and P_1_ cells have unique rates of ring assembly and initiation, we propose that the Ran pathway could govern these differences. Indeed, depletion of RAN-3 accelerates and equalizes the early phases of cytokinesis in AB and P_1_ cells. This phenotype is suppressed by co-depletion of ECT-2, supporting that this increase in kinetics occurs due to an increase in contractility. We also found that while IMB-1 (*C. elegans* importin-β) and ANI-1 are part of the Ran pathway in AB cells, IMA-3 (*C. elegans* importin-α) and IMB-1 are part of the pathway in P_1_ cells, but not ANI-1. This work highlights the importance of studying cytokinesis in different cell types and considering multiple parameters while building models of cytokinesis.

## Materials and Methods

### Strains

*C. elegans* strains (listed in the table below) were maintained according to standard protocols (Brenner, 1974) using nematode growth medium (NGM) plates seeded with *Escherichia coli* OP50.

| Strain | Genotype | Notes |
| --- | --- | --- |
| **UM463** | *cpls42[Pmex-5::mNeonGreen::PLCδ-PH::tbb-2 3'UTR; unc-119(+)] II*;<br>*ltIs37[pAA64; Ppie-1::mCherry::HIS-58; unc-119(+)] IV* | Also used to generate a tetraploid strain. |
| **OD95** | *ltIs37 [(pAA64 Ppie-1::mCherry::HIS-58 + unc-119(+))]; ltIs38 [pie-1p::GFP::PH(PLC1delta1) + unc-119(+)] III* |  |
| **SWG001** | *mex-5p::Lifeact::mKate2* |  |
| **LP162** | *cp13[nmy-2::gfp + LoxP] I*. | Also used to generate a tetraploid strain. |

### RNA interference

RNAi was carried out using feeding vectors for the induction of dsRNA expression in HT115 bacteria to target H39E23.1 (*par-1*), F28B3.8 (*imb-1*), F32E10.4 (*ima-3*), F54E7.3 (*par-3*) and T19E10.1 (*ect-2*), Y49E10.19 (*ani-1*) from the Ahringer library (Kamath et al., 2001). Strains were generously provided by Dr. Labbé (IRIC, Université de Montréal) and Dr. Roy (McGill University). Bacterial cultures were grown overnight in Luria broth (LB) with 100 µg/mL ampicillin at 37°C, then diluted 1:100 and grown at 37°C for 7 hours. The cultures were pelleted and resuspended in LB (100 µL for *ran-3, par-1* and *-3*, 300 µL for *ani-1* and *ima-3*, 400 µL for *ect-2*, 1700 µL for *imb-1*), and 50-100 µL of each resuspension was seeded onto NGM plates containing 100 µg/mL ampicillin and 1 mM IPTG. After being left to dry, 10-15 L4 hermaphrodites were placed onto each plate for 3 (*imb-1* RNAi), 24 (*par-1, par-3, ran-3, ect-2, ani-1*) or 30 hours (*ima-3* RNAi). A similar approach was used with the W02A2.6 clone (*rec-8*) to generate tetraploid worms, as described by Clarke *et al*. (Clarke et al., 2018). Specifically, L4 stage hermaphrodites of the desired strain were placed on *rec-8* RNAi plates for 8-9 days at 15°C. Then, 20 L4 stage hermaphrodites were transferred to freshly induced *rec-8* RNAi plates. After another 7-9 days, hermaphrodites that appeared longer than control were individually transferred onto OP50 plates, and maintained for successive generations by repeatedly selecting long worms. Embryos were confirmed to have higher ploidy by cell size, and staining chromosomes in fixed embryos.

### Microscopy

*C. elegans* embryos were prepared for imaging using a standard stereomicroscope by dissecting gravid hermaphrodites in M9 and transferring embryos onto a freshly prepared 2% agarose pad (Evans, 2006). Images of embryos were acquired with the 100x/1.45 NA objective on an inverted Nikon Eclipse Ti microscope fitted with a Livescan Sweptfield scanner (Nikon), Piezo Z stage (Prior), Andor IXON 897 EMCCD camera, and 488 nm and 561 nm lasers, using NIS-Elements (version 4.0, Nikon) software. Central z-planes of 0.7 μm for a total z-stack of 4 μm were collected at 5 second intervals for kymograph analysis, while 0.5 μm z-planes for a total stack of 20 μm were collected at 20 seconds to measure myosin or actin midplane cortical levels, and ring closure symmetry. All images were saved as ND2 files.

Highly inclined and laminated optical sheet (HILO) microscopy, a modified form of Total Internal Reflection Fluorescence (TIRF) (Tokunaga et al., 2008), was used to image the cortex of the AB and P_1_ cells in embryos. Embryos were transferred to agarose pads, as described above. A sub-critical incidence angle was used and adjusted until clear images of the cortex were obtained. Images were acquired with a 100x/1.49NA CFI Apo TIRF objective on an inverted Nikon Eclipse Ti microscope fitted with a TIRF arm, Photometrics Prime BSI (sCMOS) camera and 488 nm laser using NIS Elements (version 4.0, Nikon) software. Z-planes of 0.2 μm for a z-stack of 0.6 μm were collected at 2 second intervals. Images were saved as ND2 files.

### Image analysis

Only cells that successfully completed cytokinesis and had proper DNA segregation with no gross morphological defects were used for analysis, with the exception of a subset of *ect-2* (*RNAi)* AB or P_1_ cells that failed cytokinesis as described in the text. All raw data ND2 files were processed and/or analyzed in FIJI (version 2.1, NIH).

To determine the kinetics of ring closure, we used a custom macro written for FIJI to generate kymographs. Time-lapse images were staged to anaphase onset based on chromosome position (mCherry-tagged histone imaged via the 561 laser), then the change in membrane position (mNeonGreen or GFP-tagged, imaged via the 488 laser) was analyzed over time. To generate kymographs, a line with a width of 5 pixels was drawn manually over the furrow region at every time point until closure. Then, the distance between the two sides of the membrane was measured at each timepoint using the straight-line tool, and measurements were exported to Excel. The distance between the two sides at anaphase onset was set to a maximum value (100%) and used to normalize the distance throughout ingression. All n’s were averaged for each timepoint, and plotted as a function of time in seconds. Since the closure times were variable among cells, measurements were terminated when at least three cells had completed cytokinesis.

Measurements of the accumulation of actin and myosin at the midplane were performed on z-stack sum projections. A line was manually drawn along the cortex from the anterior to the posterior pole of the membrane, then 100 evenly-spaced 10 pixel-diameter circles were seeded along that line. The sum intensity for each circle was measured. To align the scans for each cell, a straight line was drawn in plane with the furrow and used to determine the linescan values located in the furrow. Each linescan within a dataset was then aligned by their furrow datapoints. Mean values were calculated for each location and positions with fewer than 3 n were not included.

HILO images were falsely colored using the mpl-inferno LUT macro in FIJI to visualize differences in myosin intensity. Cool colours (violet, dark red) reflect weaker levels compared to brighter, warmer colours (orange, yellow). The “Zoom in Images and Stacks” FIJI macro tool coded by Gilles Carpentier (Universite Paris Est Creteil Val de Marne) was used to generate images with the zoom inset (Carpentier, 2010).

### Quantitative data analysis

To measure the duration of the different phases of ring closure, graphs were analyzed using GraphPad Prism. A sigmoidal line of best fit was plotted using the averaged data for control AB and P_1_ cells, then the second derivative of the best fit line with second order smoothing (4 neighbors averaged) was plotted. The minimum and maximum x values (in seconds) of this second derivative curve represent the time points where there is a change of slope. The y value (% change in ring diameter) at the last time point of Phase 1 (ring assembly), 2 (furrow initiation) and 3 (ring constriction) was noted for each control cell. These values were used as a cut-off to define phase transitions in individual cells of control and RNAi-treated embryos. Similarly, the second derivative of the sigmoidal line of best fit for averaged tetraploid AB and P_1_ cell ingression curves was used to determine the phase transitions in tetraploid cell divisions. The phase duration for individual cells and their average was then plotted using GraphPad Prism.

To determine ring closure symmetry, the position and size of the ring were manually extrapolated for each timepoint of division (from 0% to maximum visible closure), temporally aligned, averaged and plotted. Briefly, straight lines were drawn in FIJI from one side of the ring to the other in each timepoint, then rotated to align the long axis horizontally. A 250 x 50-pixel box was then drawn around this line. These regions were rotated in Python 3 using SciKit Image (version 0.16.2) to produce an XZ view, and a projection was performed in FIJI to produce an image stack of the membrane. The ellipse tool was used to draw ellipses that matched the outline of the cell, and ellipse coordinates were recorded. In Python 3, a best-fit circle was plotted for each timepoint and each embryo. Coordinates were normalized to the first timepoint, where the center of the ring was at 0,0, and the radius at 1. The best-fit circle was averaged across all embryos within a group and plotted using the Jet colormap. To calculate symmetry, the ratio of the final ring position compared to the initial position was determined in both the x and y-axes, with >0.2 being defined as asymmetric.

To determine the orientation of myosin filaments in a defined region of the furrow of dividing AB and P_1_ cells, we used the Directionality plugin for FIJI. A region of the furrow was selected, and the plugin was run using the local gradient orientation method with 90 bins and a histogram from 0° and 90°. The plugin reports the frequency of filaments at a given angle, and fits a Gaussian function based on the highest peak in the histogram. The frequency values and Gaussian fit were plotted as a histogram in Excel. To have the center of the Gaussian fit be defined as straight (0°; perpendicular to the furrow), we subtracted the peak Gaussian value from 0°. The proportion of filaments (sum of raw histogram values) that were within the center ± two standard deviations of the Gaussian fit (referred to as the ‘Amount’) were considered to be well-aligned. The change in the proportion of well-aligned filaments was calculated by subtracting the value from control.

All graphs except those for ring closure were plotted in GraphPad Prism (version 8.4.3) and Excel (version 16.40). Ring closure symmetry graphs were plotted using Python 3 (version 3.7.6), and MatplotLib (version 3.1.3). The full code for the ring closure and membrane accumulation analyses is located at http://github.com/cmci. Graphs showing mean values are displayed with SEM bars (indicated in figure legends), and all n’s are reported in the figure legends. All figures were prepared in Adobe Illustrator.

### Pull down assays and western blots

We purified recombinant ANI-1 protein to pull down importin-β from HeLa cell lysates. To do this we cloned the RBD + C2 from ANI-1 (708-1028) into pGEX4T and pMal vectors for protein expression. We also introduced the NLS mutations K938E; K940E and K947A; K949A into the pMal:ANI-1 vector by site-directed mutagenesis. The constructs as well as control empty vectors were transformed in *E. coli* BL21 cells, grown to an ideal OD and induced with IPTG as per manufacturer’s instructions at 25°C (Sigma Aldrich for pGEX and New England Biolabs for pMal). Cells were resuspended in lysis buffer [2.5 mM MgCl2, 50 mM Tris, 150 mM NaCl, pH 7.5, 0.5% Triton X-100, 1 mM dithiothreitol (DTT), 1 mM phenylmethanesulfonyl fluoride (PMSF) and 1 X protease inhibitors; Roche], then incubated with 1 mg/mL lysozyme on ice for 30 minutes and sonicated. After sonication and centrifugation, protein was purified by incubating with glutathione agarose (GST; Sigma Aldrich) or amylose resin (MBP; New England Biolabs) for 5 hours at 4°C. The protein-bound beads were washed and stored as a. 50% slurry at 4°C. Protein concentration was measured by running samples on SDS-PAGE stained with coomassie brilliant blue and measured by densitometry against a standard curve of known BSA concentrations.

HeLa cells were cultured and transfected with a construct expressing Myc-tagged importin-β as previously described in Beaudet et al. (2017). Cells were washed then lysed on ice in lysis buffer (50 mM Tris, pH 7.6, 150 mM NaCl, 5 mM MgCl_2_, 0.5% Triton X-100, 1 mM DTT, and 1 mM PMSF). To pull down Myc:importin-β, cell lysate was incubated at 4°C with 5–10 μg of purified GST or MBP-tagged ANI-1 protein bound to glutathione or amylose beads. The beads were washed several times with wash buffer (50 mM Tris, pH 7.6, 150 mM NaCl, and 5 mM MgCl_2_), after which sample buffer was added. Samples were run by SDS–PAGE, then wet-transferred to nitrocellulose for western blotting. Transfer efficiency was visualized by ponceau S staining. Membranes were incubated with mouse anti-Myc antibodies (clone 9E10; DSHB) for 2 hours, and then washed and incubated with anti-mouse Alexa 488 secondary antibodies (Invitrogen) for 1-2 hours. After washing, blots were imaged using the 488 nm laser on a Typhoon Trio phosphoimager (GE), and the resulting images were converted to 8-bit using FIJI. Figures were prepared using Adobe Photoshop and Adobe Illustrator.

### Statistical analysis

Statistical significance was determined using unpaired Student’s *t* test with Welch’s correction on Graphpad Prism (version 8.4.3). Statistical significance was defined as: p ≥ 0.05 not significant (ns); *p = 0.01 to 0.05, **p = 0.001 to 0.01, ***p = 0.0001 to 0.001, ****p < 0.0001. An F-test was used to compare variance.

## RESULTS

### Cytokinesis occurs differently in AB and P_**1**_ **cells**

We hypothesized that cytokinesis could be differently regulated depending on cell fate and size. To test this, we studied cytokinesis of AB and P_1_ cells in the early *C. elegans* embryo, which have different fates and sizes. Since cytokinesis has not been studied extensively in these cells before, we first characterized their ring closure kinetics. To do this, we imaged embryos co-expressing GFP::PLCδ^PH^ or mNeonGreen::PLCδ^PH^ and mCherry::HIS-58 to visualize the membrane and chromatin, respectively, from anaphase onset until furrow closure with high temporal resolution **(Figure 1A)**. Kymographs produced from the images were used to measure the change in diameter until the end of ingression **(Figure S1A)**. Ring closure was not linear in AB or P_1_ cells, and we calculated the inflection points to define the three stages of ring closure; assembly, initiation, and constriction **(Figure 1B,C)**. Our results showed that while AB cells had short ring assembly and furrow initiation phases, these phases were prolonged in P_1_ cells. We also compared the duration of these phases in individual cells **(Figure 1C)**. While there was variability in both cell types, a population of P_1_ cells took longer for both phases compared to AB cells. Next, we compared the duration of ring assembly with cell size. Using simple linear regression, we found that there was no correlation between ring assembly and cell size in AB cells, while P_1_ cells had a negative correlation, suggesting that rings take longer to assemble in the smaller P_1_ cells **(Figure 1D)**.

**Figure 1.**
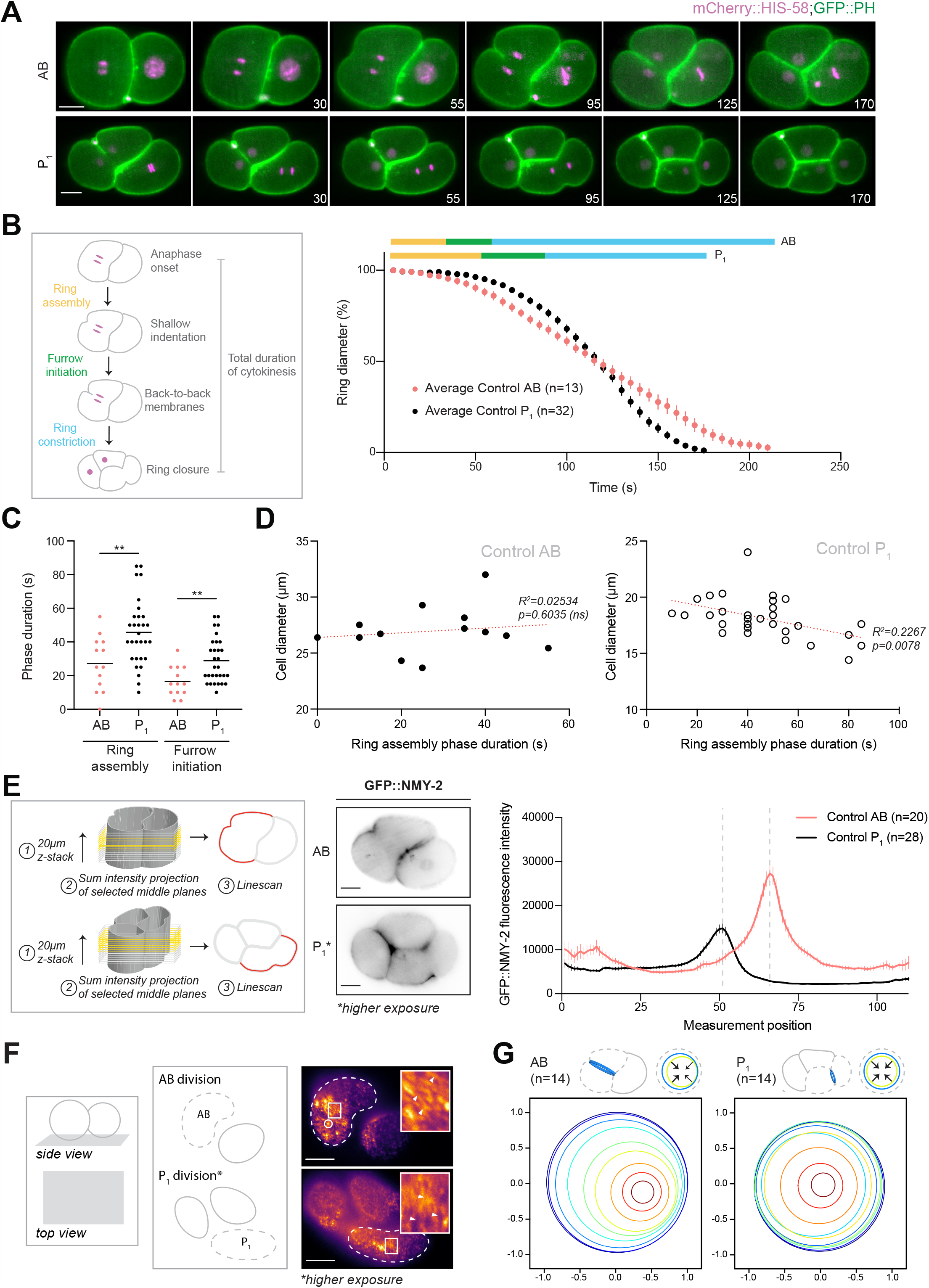
AB and P_1_ cells have unique ring closure kinetics. A) Timelapse images show furrow ingression in AB and P_1_ cells in embryos expressing mCherry::HIS-58 (magenta) and GFP::PLCδ^PH^ (green). B) Cartoon schematics show the phases of ring closure. A line graph shows ring closure (ring diameter %) in AB (coral) and P_1_ (black) cells over time. Above, bars show the durations of ring assembly (yellow), furrow initiation (green) and ring constriction (blue) phases. C) A plot shows the duration of ring assembly and furrow initiation in individual cells (average indicated by lines; **, *p*<0.01). D) Graphs show the correlation between the duration of ring assembly and diameter. The red lines show simple linear regression (R^2^ and *p* are shown). E) Schematics show how GFP::NMY-2 levels were measured. Inverted images show myosin localization in AB and P_1_ cells at furrow initiation. A graph shows GFP::NMY-2 accumulation at the midplane cortex of AB (coral) and P_1_ (black) cells. Furrow position is indicated by dotted grey lines. F) Cartoon schematics show the planes visualized by HILO imaging (AB cells at top, and P_1_ cells at bottom; dashed lines). Pseudocolored HILO images show GFP::NMY-2 in AB and P_1_ cells. The circle outlines a myosin cluster. In the zoomed inset, arrowheads point to myosin filaments. G) The cartoons show end-on ring closure. Ring closure is shown over time, with each timepoint as a different color. The x and y-axis indicate ratios of the distance from the starting position (0). For all graphs, n’s are indicated, and error bars show SEM. All scale bars are 10 µm.

We determined if the different kinetics between AB and P_1_ cells are caused by differences in the levels of cortical myosin or actin. To do this, we imaged AB and P_1_ cells expressing GFP::NMY-2 or LifeAct::mKate2. Using the central slices of a 20 µm z-stack, we drew linescans to measure myosin or actin levels along the midplane cortex **(Figure 1E, left)**. Myosin and actin localized with a bell-like distribution in AB cells, with peak levels at the furrow that were higher compared to P_1_ cells **(Figure 1E; Figure S1B)**. In P_1_ cells, myosin and actin levels were higher along the anterior cortex compared to the posterior **(Figure 1E; Figure S1B)**. Thus, myosin and actin localization are unique to each cell type.

We also compared how myosin filaments are organized in AB and P_1_ cells, as this could influence ring kinetics. We used HILO (high inclined and laminated optical sheet) microscopy to visualize GFP::NMY-2 at the cortex **(Figure 1F)**. As reported previously, we observed an asymmetric, rotational wave of myosin in AB cells **(Figure 1F, top; Movie 1)** (Singh & Pohl, 2014). We also saw clusters of myosin flowing towards the equatorial zone, resembling those seen during pseudocleavage (Munro et al., 2004; Tse et al., 2012). In contrast, there were no clusters or cortical flow in P_1_ cells **(Figure 1F, bottom; Movie 2)**, as also observed by Pimpale et al. (2020).

Next, we measured the symmetry of ring closure in AB and P_1_ cells. We expected to see asymmetric ingression in AB, but not P_1_ cells, because of the stronger cortical flows. To measure symmetry, we created a metric based on how the ring shifts in the x and y-axis relative to the starting position. When the ring closed in the middle, they were considered to be more symmetric (closer to 0) versus near the periphery (closer to 1) (**Figure S1C**). As expected, ring closure was more asymmetric in AB versus P_1_ cells **(Figure 1G; Figure S1C)**.

### Contractility controls the rate of ring assembly

We predict that the differences in myosin levels and organization in AB and P_1_ cells are controlled by differences in active RhoA, which is required for actin polymerization and myosin bipolar filament assembly **(Figure 2A)**. To test this, we mildly depleted *ect-2* without causing cytokinesis failure. The early phase of ring assembly was delayed in *ect-2(RNAi)* P_1_ cells, but comparable to control in AB cells, while the subsequent phases of furrow initiation and ring constriction were slower **(Figure 2B,C; Figure S2B)**. Additionally, there was no correlation between cell size and the duration of ring assembly in *ect-2(RNAi)* AB or P_1_ cells **(Figure 2D)**. Next, to correlate changes in myosin localization with cytokinesis phenotypes, we measured myosin levels along the midplane of *ect-2(RNAi)* embryos. There were narrower peaks of myosin in both AB and P_1_ cells compared to their control counterparts, and it was no longer asymmetrically distributed in P_1_ cells **(Figure 2E)**. The average peak of myosin in AB cells was 56% of control levels, but similar between control and *ect-2(RNAi)* P_1_ cells. Therefore, the levels of equatorial myosin are above the threshold needed to support cytokinesis in AB cells, while P_1_ cells already operate at this threshold. To validate this, we measured myosin in *ect-2(RNAi)* embryos where cells failed cytokinesis **(Figure 2F; Movie 3)**. In AB cells, peak levels were comparable to those that completed cytokinesis but myosin was not as tightly distributed, suggesting a change in its organization, while peak levels were 30% lower in P_1_ cells.

**Figure 2.**
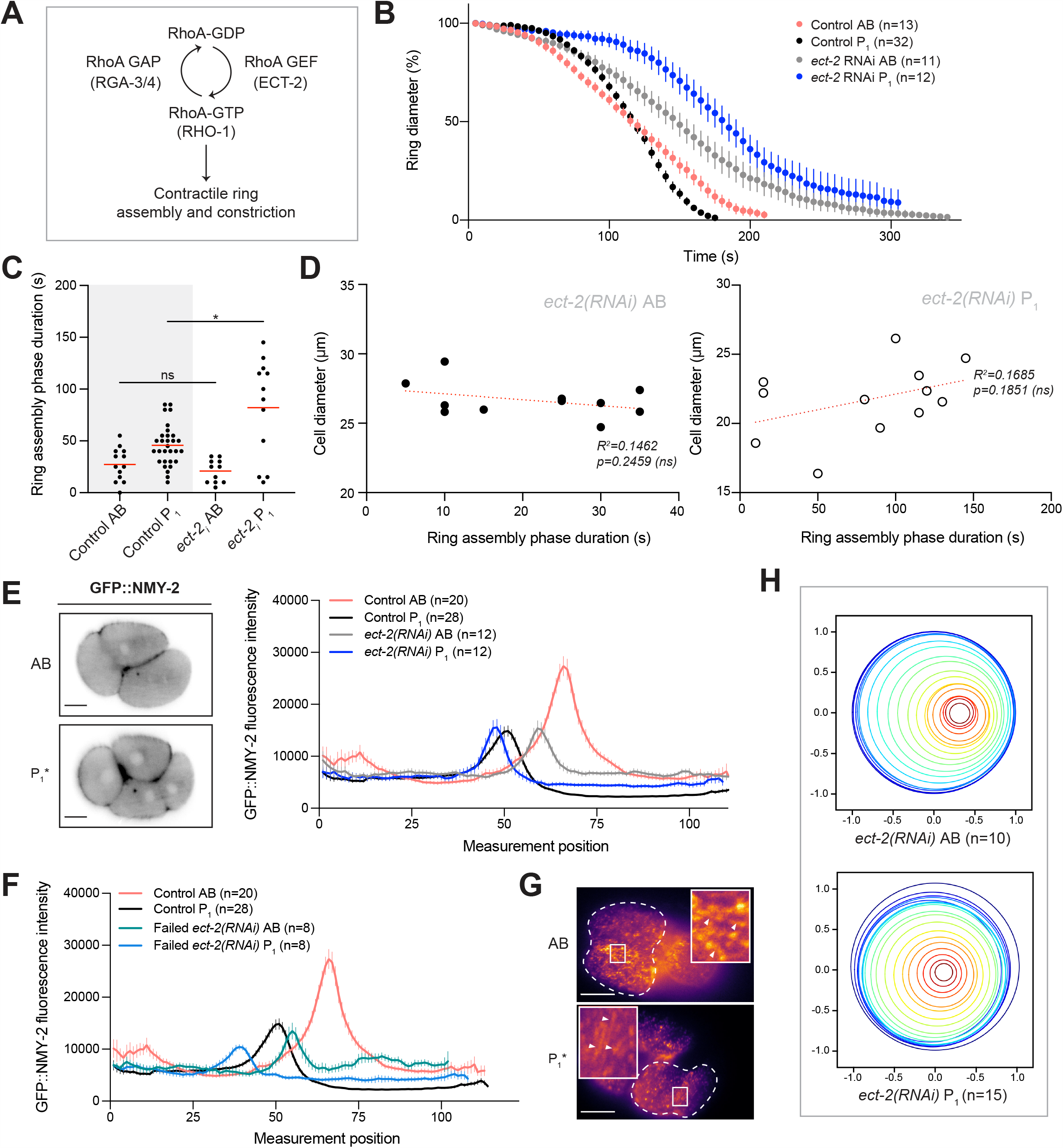
The rates of ring assembly are determined by contractility. A) A pathway shows the regulation of RhoA activity for contractile ring assembly. B) A graph shows ring closure in *ect-2(RNAi)* AB (grey) and P_1_ (blue) cells compared to control (AB, coral; P_1_, black). C) A plot shows the duration of ring assembly for individual cells (average shown by lines; *, *p*<0.05; ns, not significant). D) Graphs show the correlation between the duration of ring assembly and diameter for AB and P_1_ cells. The red lines show simple linear regression (R^2^ and *p* are shown). E) Images show GFP::NMY-2 localization in *ect-2(RNAi)* AB and P_1_ cells. A graph shows GFP::NMY-2 accumulation at the midplane cortex of control and *ect-2(RNAi)* AB (coral and grey, respectively) and P_1_ (black and blue, respectively) cells that complete cytokinesis. F) A graph shows GFP::NMY-2 accumulation in *ect-2(RNAi)* AB (green) and P_1_ (blue) cells that fail cytokinesis compared to control cells (AB; coral and P_1_; black). G) Pseudocolored HILO images show GFP::NMY-2 in a dividing *ect-2(RNAi)* AB (top) and P_1_ (bottom) cell (outlined by dashed line). In the zoomed inset, arrowheads point to myosin filaments. H) Ring closure is shown over time, with each timepoint as a different color. The x and y-axis indicate ratios of the distance from the starting position (0). For all graphs, n’s are indicated, and error bars show SEM. All scale bars are 10 µm.

To assess changes in myosin organization, we performed HILO imaging of myosin in *ect-2(RNAi)* AB and P_1_ cells **(Figure 2G; Movie 4)**. There was a dramatic loss in clusters and decreased rotational flow in AB cells compared to control cells **(Figure 2G)**. We also observed that myosin had a more punctate pattern with fewer filaments compared to control cells **(Figure 2G)**. Since flows could contribute to the alignment of myosin filaments in the ring, we determined the frequency of individual filaments in a defined region in the furrow with angles where 0° reflects a filament that is aligned with the ring, and numbers that deviate from this are less well-aligned **(Figure S3A)**. We also determined the total amount of filaments within two standard deviations of the central peak. While there was a high frequency of aligned filaments in both control AB and P_1_ cells, there was a higher total amount within two standard deviations in AB cells (‘amount’; **Figure S3A)**. In *ect-2(RNAi)* AB and P_1_ cells, we observed an increase in the frequency of filaments with angles that deviate from 0°, showing that there is a decrease in the proportion of well-aligned myosin filaments (compare **Figure 2G** with **Figure 1F; Figure S3B)**. Thus, the poor alignment of filaments in *ect2(RNAi)* AB cells could reflect a loss of cortical flows. Additionally, while ring closure remained symmetric in *ect-2(RNAi)* P_1_ cells, it was less asymmetric in AB compared to control cells **(Figure 2H; Figure S1C)**.

### Cell fate determines the rate of ring assembly

The different requirements for myosin in regulating ring closure could be attributed to fate differences between AB and P_1_ cells. For example, differences in actin and myosin in AB and P_1_ cells are likely inherited by the asymmetric P_0_ division. To test this, we assessed how disrupting fate affects ring closure by *par-1(RNAi)* and *par-3(RNAi)*. As expected, P_0_ daughter cells were equal in size and divided synchronously **(Figure 3A; Figure S2C)** (Kemphues et al., 1988). The loss of posterior PAR-1 should lead to an increase in breadth of the anterior complex, causing higher cortical contractility and AB-like kinetics, while the loss of anterior PAR-3 should cause lower contractility and P_1_-like kinetics (Cowan & Hyman, 2007; Munro et al., 2004). Surprisingly, *par-3(RNAi)* and *par-1*(*RNAi*) P_0_ daughter cells had kinetics that were similar to AB cells, and even faster, respectively (**Figure 3B,C**). In line with their AB-like (or faster) kinetics, we found no correlation between cell size and the duration of ring assembly **(Figure 3D)**.

**Figure 3.**
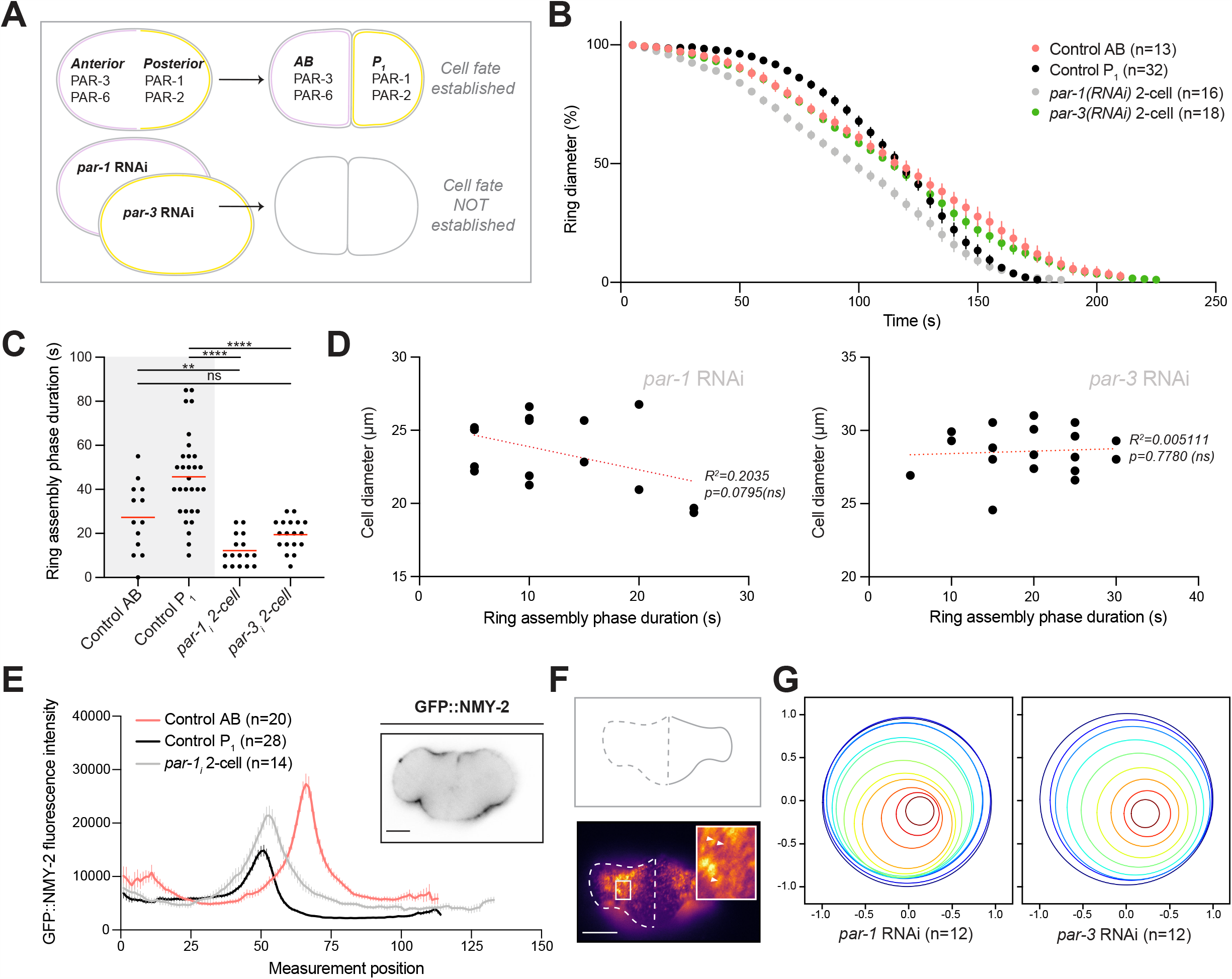
Cell fate impacts cytokinesis in the early embryo. A) A cartoon shows the distribution of PAR proteins in the P_0_ zygote, and how their depletion disrupts cell fate. B) A graph shows ring closure in *par-1(RNAi)* (grey) and *par-3(RNAi)* (green) P_0_ daughters compared to control cells (AB, coral; P_1_, black). C) A plot shows the duration of ring assembly for individual cells (average shown by lines; **, *p*<0.01; ****, *p*<0.0001; ns, not significant). D) Graphs show the correlation between the duration of ring assembly and diameter for *par-1* and *par-3(RNAi)* P_0_ daughter cells. The red lines show simple linear regression (R^2^ and *p* are shown). E) A graph shows GFP::NMY-2 accumulation at the midplane cortex of control AB (coral) and P_1_ (black) cells, and *par-1(RNAi)* (grey) P_0_ daughter cells (GFP::NMY-2 localization shown in inset). F) Pseudocolored HILO images show GFP::NMY-2 in a dividing *par-1(RNAi)* P_0_ daughter cell. In the zoomed inset, arrowheads point to myosin filaments. G) Ring closure is shown over time, with each timepoint as a different color. The x and y-axis indicate ratios of the distance from the starting position (0). For all graphs, n’s are indicated, and error bars show SEM. All scale bars are 10 µm.

Next, we determined if myosin levels and organization support the rapid kinetics in *par-1(RNAi)* cells. Peak myosin levels were higher than in control P_1_ cells, but not as high as control AB cells (78%) **(Figure 3E)**. Since myosin was distributed equally between the daughter cells and localized over a broader area, this could dilute the pool in a single location. HILO imaging revealed strong cortical flows and broad swaths of densely packed filaments in both cells **(Figure 3F; Movie 5)**. The filaments also appeared to be well-aligned in the furrow of *par-1(RNAi)* cells **(Figure 3F; Movie 5)**. Strong cortical flows could facilitate the localization and alignment of myosin filaments to support their enhanced kinetics. Indeed, ring closure occurred asymmetrically in *par-3(RNAi)* and *par-1(RNAi)* P_0_ daughter cells **(Figure 3G; Figure S1C)**.

### Cell size influences the rate of ring assembly

We next wanted to determine if ring closure kinetics are cell-size dependent, especially during the earlier phases of cytokinesis. We generated tetraploid (4n) mNeonGreen::PLCδ^PH^; mCherry::HIS-58 embryos, which have a 1.3-fold and 1.46-fold increase in AB (27 – 35 µm) and P_1_ (18.6 – 27.2 µm) cell size, respectively **(Figure 4A; Figure S2D)**. The average size of tetraploid P_1_ cells is nearly identical to diploid AB cells, and they had similar ring assembly and furrow initiation phases **(Figure 4B,C)**. Surprisingly, both phases took much longer in tetraploid AB cells compared to diploid AB or P_1_ cells **(Figure 4B,C)**. There was no correlation between the duration of ring assembly and cell size for tetraploid AB cells, and a stronger negative correlation in tetraploid P_1_ cells **(Figure 4D)**. The negative correlation in tetraploid P_1_ cells was surprising given that their kinetics resembled AB cells. Several factors could contribute to this correlation, including the Ran pathway (see next section), which increases in strength with ploidy (Hasegawa et al., 2013).

**Figure 4.**
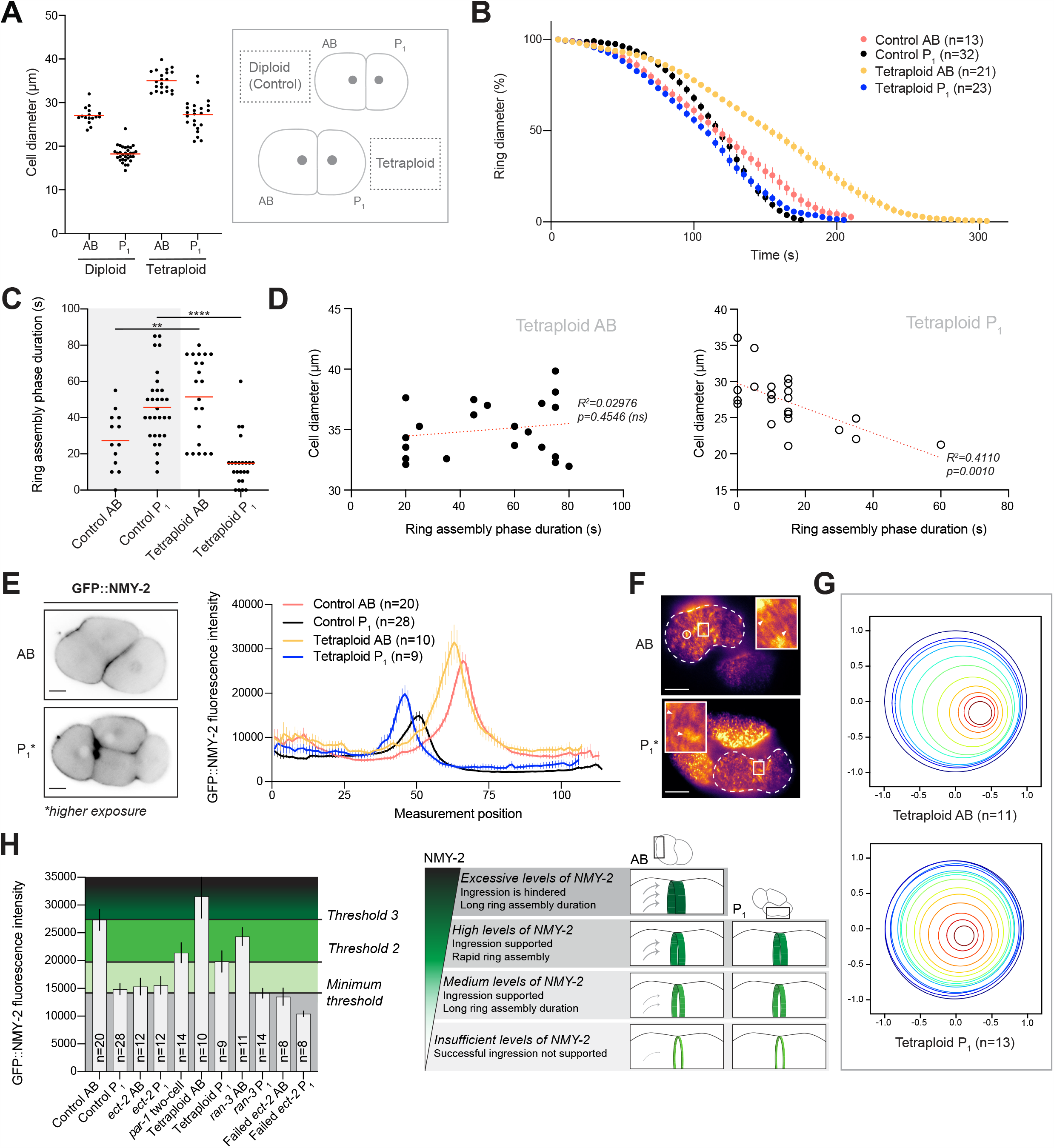
Cell size impacts AB and P_1_ cell cytokinesis. A) A graph shows the size of diploid and tetraploid AB and P_1_ cells. Cartoons highlight their relative sizes. B) A graph shows ring closure in tetraploid AB (yellow) and P_1_ (blue) cells compared to control cells (AB, coral; P_1_, black). C) A plot shows the duration of ring assembly for individual cells (average shown by lines; **, *p*<0.01; ****, *p*<0.0001). D) Graphs show the correlation between the duration of ring assembly and diameter. The red lines show simple linear regression (R^2^ and *p* are shown; ns is not significant). E) Images show GFP::NMY-2 localization in tetraploid AB and P_1_ cells. A graph shows GFP::NMY-2 accumulation at the midplane cortex of tetraploid AB (yellow) and P_1_ (blue) cells compared to control (AB, coral; P_1_, black). F) Pseudocolored HILO images show GFP::NMY-2 in a tetraploid AB and P_1_ cell (outlined by the dashed line). The circle shows a myosin cluster. In the zoomed inset, arrowheads point to myosin filaments. G) Ring closure is shown over time, with each timepoint as a different color. The x and y-axis indicate ratios of the distance from the starting position (0). H) The graph shows the threshold levels for myosin in ring closure in AB and P_1_ cells, based on the different treatments. A schematic shows the correlation between these thresholds and ring assembly. The contractile ring is in green, and the different myosin levels are indicated by shades of green, with arrows indicating flows. For all graphs, n’s are indicated, and error bars show SEM. All scale bars are 10 µm.

To determine if the different kinetics in tetraploid AB and P_1_ cells correlates with myosin levels and/or organization, we generated tetraploid embryos expressing GFP::NMY-2. Peak myosin levels in tetraploid AB cells were higher and more broadly distributed compared to diploid AB cells. There was also an increase in peak myosin levels in tetraploid versus diploid P_1_ cells **(Figure 5E)**. HILO imaging revealed a high number of myosin clusters in tetraploid AB cells, with densely packed filaments throughout the cortex **(Figure 4F; Movie 6)**. Tetraploid P_1_ cells also appeared to have a greater density of myosin compared to diploid P_1_ cells **(Figure 4F; Movie 6)**. Since the myosin filaments appeared to be well-aligned **(Figure 4F; Movie 6)**, the slower kinetics could be due to excessive force generation outside the furrow. Despite this, the AB cells retained sufficient flows to support asymmetric ring closure. The rings in tetraploid P_1_ cells closed symmetrically, but they were more asymmetric than in diploid cells **(Figure 4G; Figure S1C)**.

**Figure 5.**
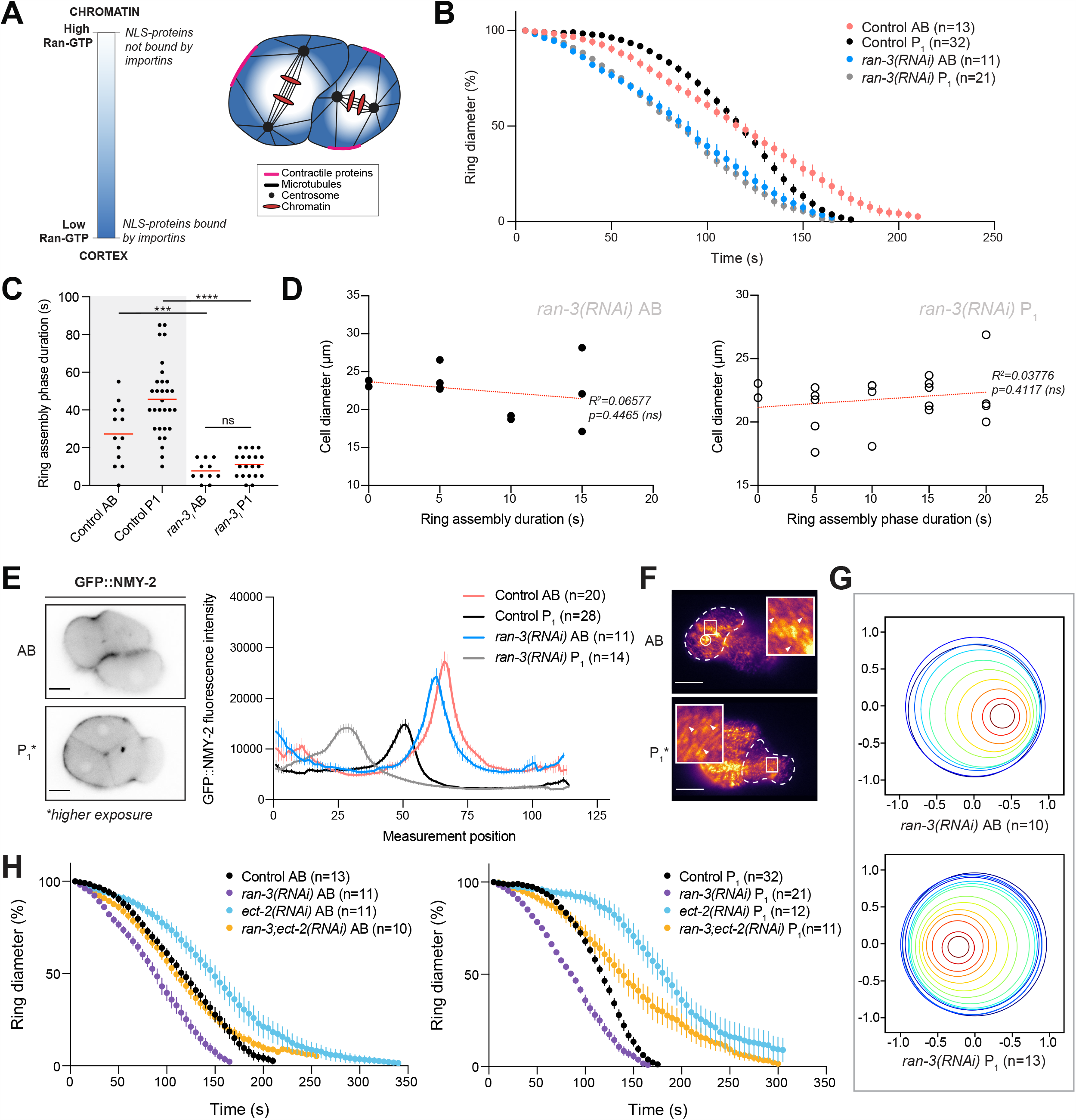
Ran regulates cytokinesis in the two-cell *C. elegans* embryo. A) A cartoon shows the importin gradient in the two-cell embryo. Ran-GTP is high (white) near chromatin (red) and low near the cortex (blue) where importins are free to bind to NLS-containing contractile proteins (pink). Centrosomes and microtubules are shown in black. B) A graph shows ring closure in *ran-3(RNAi)* AB (blue) and P_1_ (grey) cells compared to control (AB, coral; P_1_, black). C) A plot shows the duration of ring assembly for individual cells (average shown by lines; ***, *p*<0.001; ****, *p*<0.0001; ns, not significant). D) Graphs show the correlation between diameter and the duration of ring assembly. The red lines show simple linear regression (R^2^ and *p* are shown). E) Images show GFP::NMY-2 localization in *ran-3(RNAi)* AB and P_1_ cells. A graph shows GFP::NMY-2 accumulation at the midplane cortex of control and *ran-3(RNAi)* AB (coral and blue, respectively) and P_1_ (black and grey, respectively) cells compared to control (AB, coral; P_1_, black). F) Pseudocolored HILO images show GFP::NMY-2 in a dividing *ran-3(RNAi)* AB (top) and P_1_ (bottom) cell (outlined by dashed line). In the zoomed inset, arrowheads point to myosin filaments. G) Ring closure is shown over time, with each timepoint as a different color. The x and y-axis indicate ratios of the distance from the starting position (0). H) A) Graphs show ring closure in AB (left) and P_1_ (right) *ran-3* (purple), *ect-2(RNAi)* (blue) and *ran-3; ect-2(RNAi)* (yellow) cells compared to control (black). For all graphs, n’s are indicated, and error bars show SEM. All scale bars are 10 µm.

Our results suggest that there are distinct thresholds for myosin in AB and P_1_ cells. We found that the rings in AB cells ingress asymmetrically and have more filamentous myosin than P_1_ cells, which supports an ideal threshold for rapid ring assembly **(Figure 4H)**. The rings in P_1_ cells ingresses symmetrically, and already operate near or at the minimum required threshold for myosin **(Figure 4H)**. Prolonged ring assembly in these cells may ensure that they divide asymmetrically to produce the correct founder cells in future divisions. Our *ect-2* RNAi data revealed the minimum threshold required to support ring assembly in AB and P_1_ cells **(Figure 4H)**. Conversely, our tetraploid data revealed the maximum threshold **(Figure 4H)**. The range for an ideal myosin threshold for efficient ring assembly is between those observed for control AB and tetraploid P_1_ cells, or P_0_ daughters after depletion of PAR-3 or PAR-1. Thus, our data shows that contractility, cell fate and size are factors that contribute to the unique kinetics observed in AB and P_1_ cells. We propose that different mechanisms regulate cytokinesis in these cell types. In particular, we previously showed that the Ran pathway regulates cytokinesis in cultured human cells by sensing chromatin position (Beaudet et al., 2017; Beaudet et al., 2020), and we hypothesize that this pathway may regulate cytokinesis differently in AB vs. P_1_ cells.

### The Ran pathway regulates cytokinesis in AB and P_1_ cells

We determined if the Ran pathway regulates the early phases of ring closure in AB and P_1_ cells. Our model is that importin-binding to NLS-containing cortical proteins could increase their enrichment at the equatorial cortex. We previously showed in human cells that anillin requires binding to importin-beta for its localization and function during cytokinesis (Beaudet et al., 2017; Beaudet et al., 2020). To test this, we partially depleted RAN-3 (RCC1) to decrease the levels of Ran-GTP and increase the pool of importins that can bind to NLS-containing proteins **(Figure 5A)**. Ring assembly was faster in both AB and P_1_ cells after *ran-3(RNAi)*, and kinetics were equalized in the two cell types (**Figure 5B; Figure S2D**). Individual cells were highly clustered and there was no significant difference in the duration of ring assembly for AB compared to P_1_ cells **(Figure 5C)**. To ensure that this change in kinetics was not due to a change in polarity, we imaged embryos co-expressing GFP::PH with PGL-1::RFP, which is a marker of P granules **(Figure S4A)**. We saw that P granules segregated asymmetrically to P_1_ cells in embryos after *ran-3(RNAi)*, similar to control embryos (Strome & Wood, 1982). We also saw no correlation between cell size and the duration of ring assembly in either *ran-3(RNAi)* AB or P_1_ cells **(Figure 5D)**.

Next, we determined if the rapid ring assembly kinetics in *ran-3(RNAi)* cells was due to an increase in myosin levels and/or organization. Peak myosin levels in *ran-3(RNAi)* AB cells were similar but slightly lower than control cells, while P_1_ cells had similar peak levels but a broader distribution of myosin, similar to *par-1* RNAi embryos **(Figure 5E)**. HILO imaging revealed that both AB and P_1_ cells had densely packed filamentous myosin at their cortex, and AB cells had strong cortical flows **(Figure 5F; Movies 7 and 8)**. In support of their rapid kinetics, the myosin filaments appeared to be well-aligned in both AB and P_1_ *ran-3(RNAi)* cells, but especially in P_1_ cells compared to control **(Figure S3)**. These results suggest that Ran-GTP appears to affect the organization of myosin at the cortex rather than the levels per se. As expected with strong cortical flows, the rings closed asymmetrically in AB cells **(Figure 5G)**. However, they also closed asymmetrically in P_1_ cells, even though there were no obvious flows **(Figure 5G)**. While it is not clear how this asymmetry is driven, there could be asymmetric forces generated outside the ring.

Next, we determined if the changes in cytokinesis in AB and P_1_ cells after *ran-3(RNAi)* were due to changes in contractility. To test this, we co-depleted ECT-2 and RAN-3, and measured ring closure kinetics in AB and P_1_ cells **(Figure 5H)**. The faster ring assembly and/or initiation phases caused by *ran-3(RNAi)* were suppressed by partial *ect-2(RNAi)* in both AB and P_1_ cells **(Figure 5H)**. Therefore, our data supports that Ran-GTP levels influence contractility in these two cell types.

### The Ran pathway functions through different components in AB and P_1_ cells

Since Ran-GTP levels regulate ring assembly and initiation kinetics in AB and P_1_ cells, we determined if Ran functions by regulating importins. To do this, we depleted IMA-3 (IMA-3; importin-α) or IMB-1 (IMB-1; importin-β). *C. elegans* has three importin-α homologues (IMA-1, −2 and −3), but IMA-1 depletion has no obvious phenotype, and IMA-2 is essential for spindle assembly precluding its use in this study (Askjaer et al., 2002; Geles & Adam, 2001). Interestingly, while *ima-3(RNAi)* caused faster ring assembly and initiation kinetics in both AB and P_1_ cells, *imb-1(RNAi)* caused faster kinetics in P_1_ cells, but not in AB cells **(Figure 6A)**. Co-depletion of *imb-1* suppressed the rapid kinetics in *ima-3(RNAi)* AB cells, while in P_1_ cells, there was only partial suppression **(Figure S4B)**. This differential response suggests that the Ran pathway functions through different importins in different cell types.

**Figure 6.**
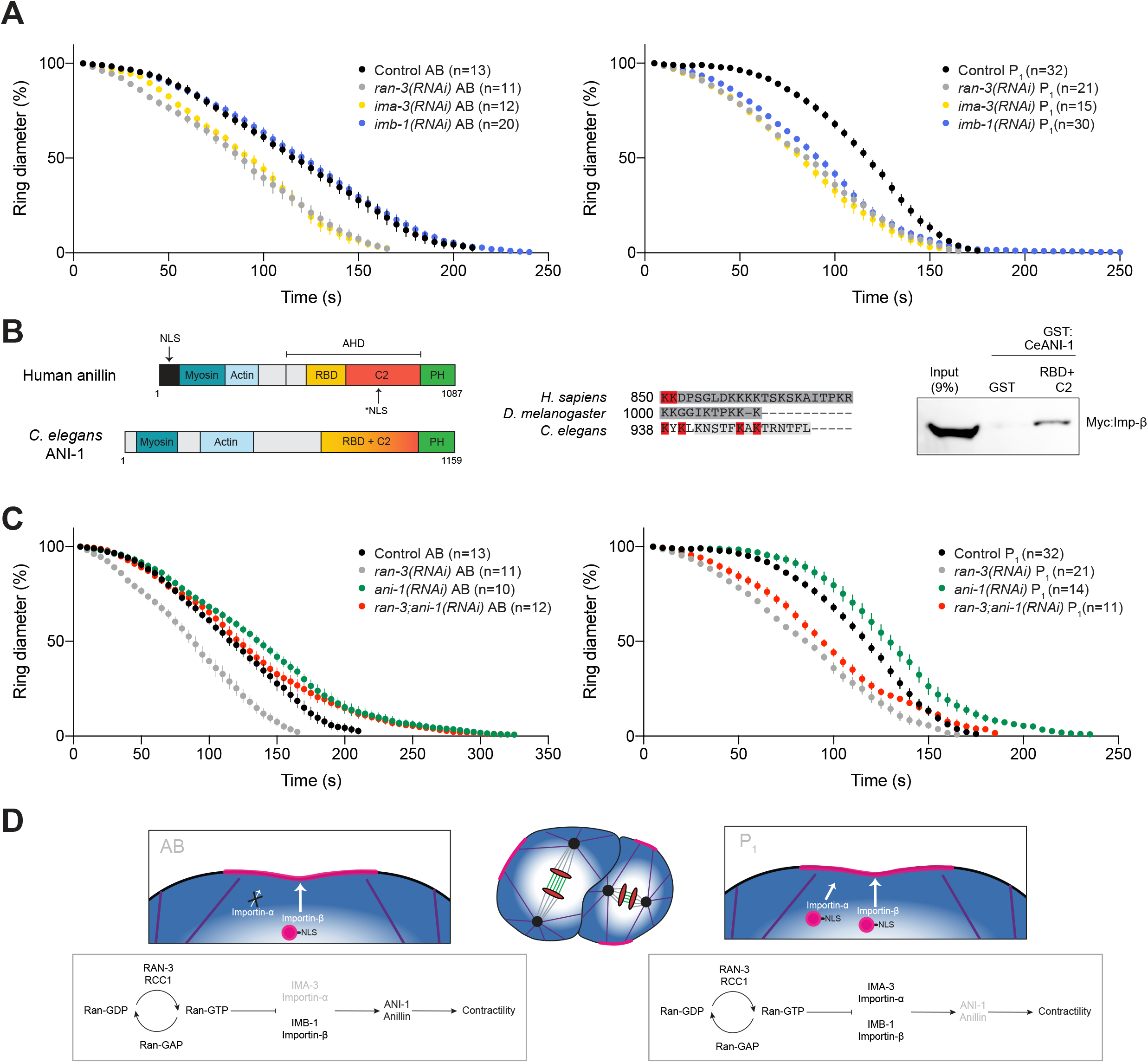
Ran regulation differs between AB and P_1_ cells. A) Graphs show ring closure in AB (left) and P_1_ (right) *ran-3(RNAi)* (grey), *ima-3(RNAi)* (yellow) and *imb-1(RNAi)* (blue) cells compared to control (AB, coral; P_1_, black). B) Cartoons show the structure of human anillin and *C. elegans* ANI-1 with the various binding domains [myosin binding domain, blue-green; F-actin binding domain, blue; RhoA-GTP binding domain (RBD), yellow; C2 domain, red; Pleckstrin Homology domain (PH), green]. In human anillin, the N-terminal NLS mediates nucleocytoplasmic transport, while the C-terminal NLS (asterisk) binds to importin-β to regulate its localization and function. The sequence of the C-terminal NLS is shown for various anillin homologues, and the residues that were mutated to disrupt importin binding are shown in red. To the right, a western blot shows Myc-tagged importin-β from HeLa cell lysates (input) and after pull-down with recombinant, purified GST or GST-tagged ANI-1 (RBD + C2). C) Graphs show ring closure in AB (left) and P_1_ (right) *ran-3(RNAi)* (grey), *ani-1(RNAi)* (green) and *ran-3; ani-1(RNAi)* (red) cells compared to control (AB, coral; P_1_, black). D) Cartoons summarize how the Ran pathway differs in AB (left) and P_1_ (right) cells. As indicated in the pathway maps, while Ran signalling regulates cytokinesis in both cell types, it does so through different targets and importins.

Next, we determined if ANI-1 (anillin) is a target of the Ran pathway in AB and P_1_ cells. Currently, anillin is the only cytokinesis protein known to be regulated by importin-β-binding for its cortical function in human cells. Most of ANI-1 shares homology with human anillin **(Figure 6B)**, and the NLS in the C2 domain appears to be conserved **(Figure 6B)**. Indeed, GST-tagged ANI-1 (RBD + C2) pulled down Myc-tagged human importin-β from cell lysates **(Figure 6B)**. This binding was reduced by point mutations in the NLS **(Figure S4C)**. Therefore, this data suggests that ANI-1 has the potential to be regulated by importin-binding and the Ran pathway in *C. elegans*

To test if ANI-1 function could be influenced by changes in Ran-GTP levels, we co-depleted ANI-1 and RAN-3. While *ani-1(RNAi)* caused only minor delays in ring assembly and/or initiation in AB and P_1_ cells, it suppressed ring closure kinetics in *ran-3(RNAi)* AB cells as expected for a component of the pathway, but not in P_1_ cells (**Figure 6C**). Thus, in addition to having unique importin requirements, our data shows that the Ran pathway controls other cortical regulators in P_1_ cells.

## DISCUSSION

### AB and P_1_ cells have distinct ring closure kinetics

We demonstrate that AB and P_1_ cells in the *C. elegans* embryo have unique ring closure kinetics, which are influenced by cell fate and size and governed differently by the Ran pathway. We found that ring assembly and initiation are faster in AB cells compared to P_1_ cells, and this correlates with higher levels of equatorial myosin and aligned filaments that appear to flow into the contractile ring **(Figure 1)**. This is consistent with prior studies suggesting that long-range flows promote ring assembly (Illukkumbura et al., 2020; Khaliullin et al., 2018; Reymann et al., 2016; Singh & Pohl, 2014). These strong cortical flows also position the midbody remnant in AB cells, which influences spindle position in P_1_ cells for proper cell fate determination (Singh & Pohl, 2014). We also found that the ring closes asymmetrically in AB, but not P_1_ cells, which could be due to these flows causing greater pull from one side of the ring (Khaliullin et al., 2018; Maddox et al., 2007). Additionally, the clusters observed on the AB cortex may facilitate the alignment and organization of actin and myosin and/or compression (Khaliullin et al., 2018; Reymann et al., 2016). In contrast, it takes longer to accumulate the threshold myosin levels needed for ring assembly in P_1_ cells, which lack cortical flows and have lower myosin levels, while constriction occurs with less hindrance due to lower cortical tension that antagonizes furrowing (Silva et al., 2016). These data indicate that the levels of myosin correlate with ring closure kinetics **(Figure 4H)**. In support of this, the different kinetics in AB and P_1_ cells were largely suppressed by partial depletion of ECT-2, which generates active RhoA for myosin activity and F-actin assembly (Green et al., 2012) **(Figure 2)**. Our findings revealed that AB and P_1_ cells have similar minimum threshold levels of myosin. While AB cells have higher levels of myosin and are sensitized to cytokinesis failure when they drop to ∼50%, P_1_ cells are more robust and already operate at the minimum threshold. Surprisingly, even though myosin filaments were poorly aligned and flows were reduced, rings still closed asymmetrically in *ect-2(RNAi)* AB cells, suggesting that multiple factors contribute to this phenomenon, which is still not fully understood.

### Differences in AB and P_1_ ring closure kinetics are fate-dependent

The higher levels of myosin in AB versus P_1_ cells reflect its unequal inheritance from the P_0_ division, implicating cell fate as a key factor that influences cytokinesis. Indeed, disrupting cell fate by depletion of PAR-1 or PAR-3 equalized ring closure kinetics between the daughter cells **(Figure 3)**. While PAR-1 is part of the posterior complex, PAR-3 is part of the anterior complex where cortical contractility is enriched (Cowan & Hyman, 2007; Rose & Gonczy, 2014). Thus, we expected that their depletion would cause different ring closure kinetics; loss of PAR-1 should cause daughters to resemble AB cells, while loss of PAR-3 should cause daughters to resemble P_1_ cells. However, we found that loss of either PAR-1 or PAR-3 caused the daughters to be more AB-like in their kinetics. Myosin accumulated to levels between AB and P_1_, but the peaks were broad and the same in both daughters. We also observed strong cortical flows with well-aligned filaments in both daughters, which ingressed asymmetrically. It is not clear why PAR-3 depletion caused kinetics that were more AB-like. Multiple factors contribute to the global cortical contractility observed in oocytes, which becomes inhibited at the posterior cortex by PAR-1/PAR-2, and expansion of PAR-1/PAR-2 complexes may be insufficient to entirely suppress these factors (Cowan & Hyman, 2007; Rose & Gonczy, 2014). In addition, this early pool of contractile myosin would be distributed equally to the daughters, instead of being asymmetrically enriched in AB cells.

### Size governs differences in AB and P_1_ ring closure kinetics

Cell size also influences ring closure kinetics in AB and P_1_ cells. We analyzed this in several ways. First, we tested the relationship between cell size and the duration of ring assembly **(Figures 1-5D)**. We found that while AB cells had no correlation in any condition, P_1_ cells showed a negative correlation in control diploid and tetraploid embryos. This suggests that it takes the ring longer to assemble in larger P_1_ cells. Since this relationship was retained in tetraploid P_1_ cells, which otherwise showed AB-like kinetics, this suggests that this regulation is fate-dependent. In support of this, we found that cells depleted of PAR-3 or PAR-1 failed to show a correlation between cell size and duration of ring assembly. The correlation in diploid and tetraploid P_1_ cells also was lost with *ect-2(RNAi)* or *ran-3(RNAi)*, both of which equalize kinetics between AB and P_1_ cells by either suppressing or enhancing contractility, respectively. We also determined how increasing the size of embryos affects ring closure kinetics **(Figure 4)**. We were surprised to observe that all of the stages of ring closure were slower in AB, and faster in tetraploid P_1_ cells compared to diploid cells. Interestingly, the size of tetraploid P_1_ cells is the same as diploid AB cells, and myosin levels were higher in both AB and P_1_ cells compared to control cells. We observed large swaths of myosin filaments and clusters in expanded regions of the cortex in AB cells, while there appeared to be an increase in aligned filaments in P_1_ cells compared to their diploid counterparts. We propose that the delayed kinetics in AB cells reveal a maximum threshold of myosin, where excess levels hinder vs. facilitate ring assembly and closure **(Figure 4H)**. The tension generated by these excess filaments could counter those that would facilitate the various stages of ring assembly and constriction.

### The Ran pathway regulates cytokinesis differently in AB and P_1_ cells

Our data shows that the mechanisms regulating cytokinesis vary with cell fate and size. Our prior studies of cytokinesis in human cells discovered a role for Ran-GTP in coordinating the position of the contractile ring with segregating chromosomes, and we proposed that this mechanism could be differently required depending on cell fate and/or size (Beaudet et al., 2017; Beaudet et al., 2020) **(Figure 5A)**. This pathway relies on a gradient of active Ran associated with chromatin, which forms inverse to a gradient of importins free to bind to NLS-containing proteins (Ozugergin & Piekny, 2020). Importantly, the Ran/importin gradient was previously shown to vary with cell size and ploidy (Deng et al., 2007; Hasegawa et al., 2013). In human cells, anillin contains an NLS that binds to importin-β and is required for its localization and function during cytokinesis (Beaudet et al., 2017; Beaudet et al., 2020). We found that this NLS is conserved in ANI-1, and we propose that binding to importin-β similarly facilitates its cortical recruitment and function during cytokinesis (**Figure 6B**). Lowering Ran-GTP levels via depletion of the Ran GEF RAN-3/RCC1 equalized ring closure kinetics in AB and P_1_ cells, which assembled rings more rapidly compared to control cells (**Figure 5**). The levels of myosin were similar to control cells, but distributed over a broader area, suggesting a change in its organization that facilitates contractility. We found that the rapid kinetics were due to an increase in myosin activity, as *ect-2(RNAi)* suppressed kinetics in both AB and P_1_ cells. We also found that the Ran pathway functions through different components in AB and P_1_ cells **(Figure 6)**. For example, depletion of IMA-3 also led to rapid kinetics in AB and P_1_ cells, while depletion of IMB-1 caused rapid kinetics only in P_1_ cells. The dogma in the field is that importin-α binds to the NLS of proteins and acts as an adaptor protein for importin-β (Ozugergin & Piekny, 2020; Xu & Massague, 2004). However, data from a number of labs suggest that importin-α and −β can bind independently to NLS-containing proteins (Ozugergin & Piekny, 2020). We propose that in P_1_ cells, either IMA-3 or IMB-1 can bind to and regulate cortical proteins (or their regulators), which can influence their localization and function in cytokinesis. However, in AB cells, IMB-1 carries out this function. We also found that the targets of the Ran pathway may differ from cell to cell. Our co-depletion experiments suggest that ANI-1 is a target of the Ran pathway in AB, but not P_1_ cells. We are currently identifying other NLS-containing cytokinesis proteins that are targeted by this pathway.

Cumulatively, our data shows that AB and P_1_ cell cytokinesis differs from the P_0_ zygote, and that ring assembly and initiation kinetics relies on several factors such as cell fate and size. We also found that the Ran pathway regulates cytokinesis differently in AB and P_1_ cells, demonstrating that the mechanisms regulating cytokinesis differs with fate. These findings highlight the need to study cytokinesis in different cell types, and to consider how microtubule-independent mechanisms influence ring kinetics.

## AUTHOR CONTRIBUTIONS

A.P. supervised the project, designed experiments and edited the manuscript. I.O. performed the majority of experiments and formal analysis, and was the primary author of the manuscript. C.L. wrote macros for FIJI and MATLAB and performed formal analysis. K.M. contributed to data collection. The authors declare no conflict of interest.

## ACKNOWLEDGMENTS

We thank members of the Piekny lab for helpful discussions and feedback, Mathew Duguay for experimental support, and Daniel Beaudet for his contribution to the ANI-1 pull-down experiment. We thank J.C. Labbé (IRIC, Université de Montréal, Montreal, QC, Canada), A.S. Maddox (UNC Chapel Hill, NC, USA), S.W. Grill (Max Planck Institute of Molecular Cell Biology and Genetics, Dresden, Germany), and the *Caenorhabditis* Genetics Center (CGC), which is funded by NIH Office of Research Infrastructure Programs (P40 OD010440), for worm strains. This work was funded by the Natural Sciences and Engineering Research Council of Canada [RGPIN-2017-04161]. We acknowledge the support of the Natural Sciences and Engineering Research Council of Canada (NSERC), [CREATE 511601-2018].

## FIGURE LEGENDS

**Figure S1.**
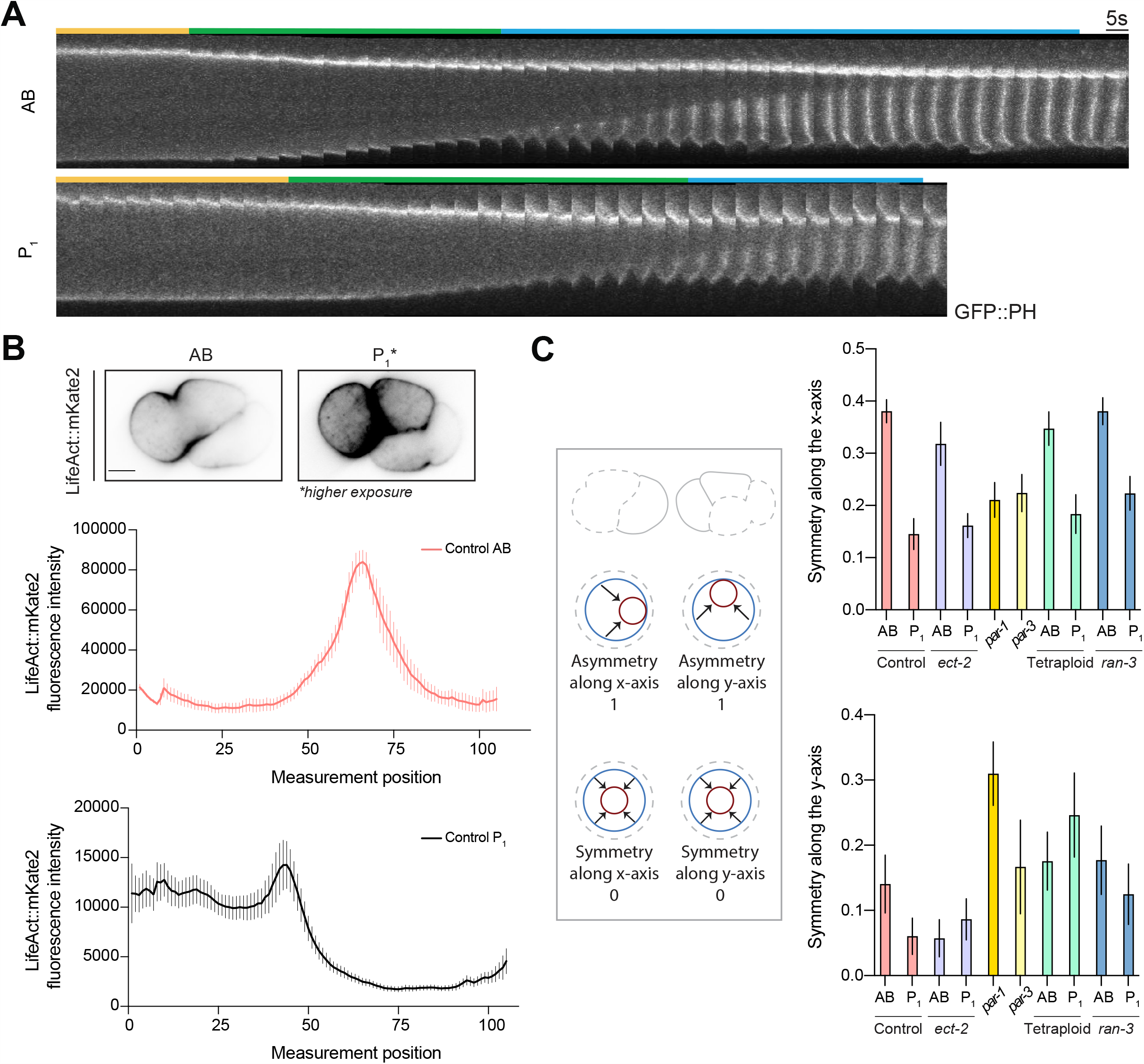
Characterizing ring closure in AB and P_1_ cells. A) Kymographs were generated from a region of interest at the furrow region in AB (top) and P_1_ (bottom) cells, from images acquired at 5-second intervals from anaphase onset until closure. The bars above each kymograph indicate the duration of ring assembly (yellow), furrow initiation (green) and ring constriction (yellow) phases. B) Images show LifeAct::mKate2 localization in control AB and P_1_ cells. Graphs show the average accumulation of LifeAct::mKate2 at the midplane cortex of AB (top) and P_1_ (bottom) cells. C) Cartoon embryos and end-on views show how the symmetry of ring closure was quantified. Asymmetric ring closure (along the x or y-axis) reflects values closer to 1, while those closest to 0 are symmetric. The graphs to the right show the values for the x-axis (top) and y-axis (bottom). All error bars show SEM. All scale bars are 10 µm.

**Figure S2.**
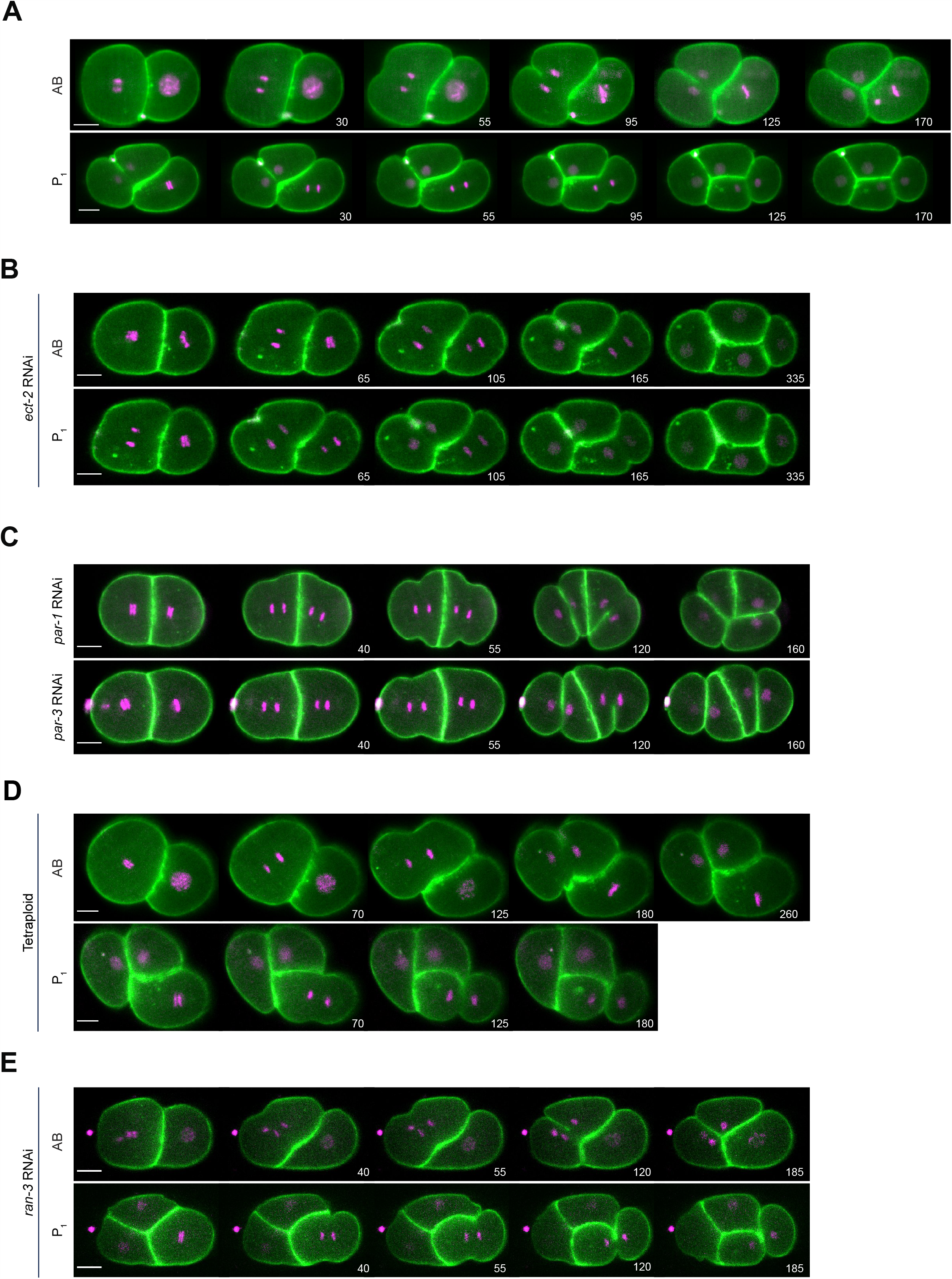
Movies of AB and P_1_ cells in cytokinesis in different conditions. Timelapse images of A) control (also shown in Figure 1A), B) *ect-2(RNAi)*, C) *par-1(RNAi)* (top) and *par-3(RNAi)* (bottom), D) tetraploid and E) *ran-3(RNAi)* embryos co-expressing mCherry::HIS-58; GFP:: PLCδ^PH^ (A, E) or mCherry::HIS-58 and mNeonGreen::PH (B-D). All scale bars are 10 µm.

**Figure S3.**
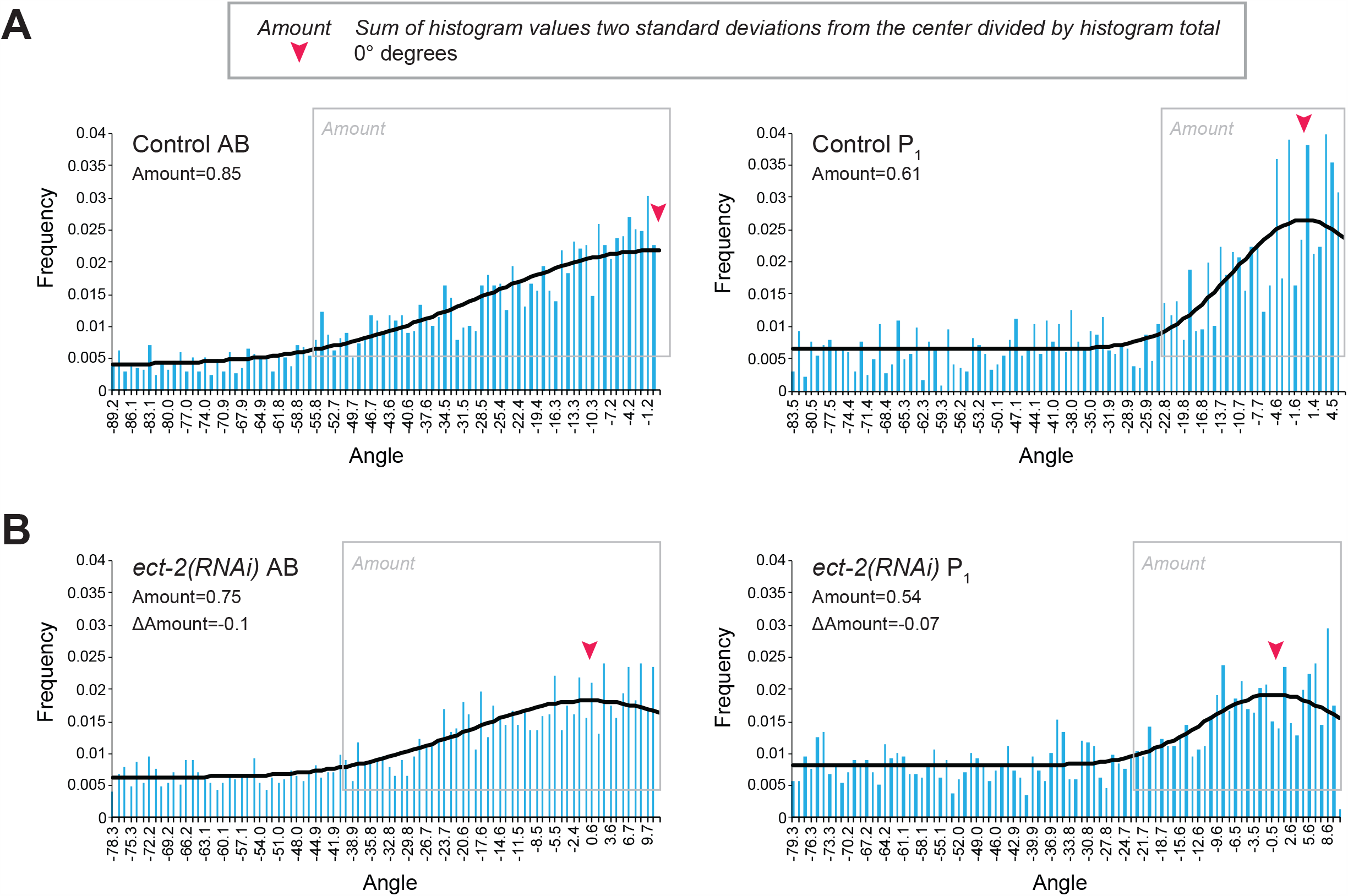
Directionality of filaments in the furrow of AB and P_1_ cells under different conditions. Histograms show the frequency distribution of myosin filaments at the angles indicated on the x-axis. Measurements were taken in the furrow of A) control and B) *ect-2(RNAi)*, cells shown in Figures 1F and 2G, respectively. Filaments that were considered to be well-aligned (center ± two standard deviations) are indicated by the grey boxes, and the total proportion of filaments in this region are indicated as ‘Amount’ in each graph. 0° (straight) is indicated by the red arrowhead in each graph.

**Figure S4.**
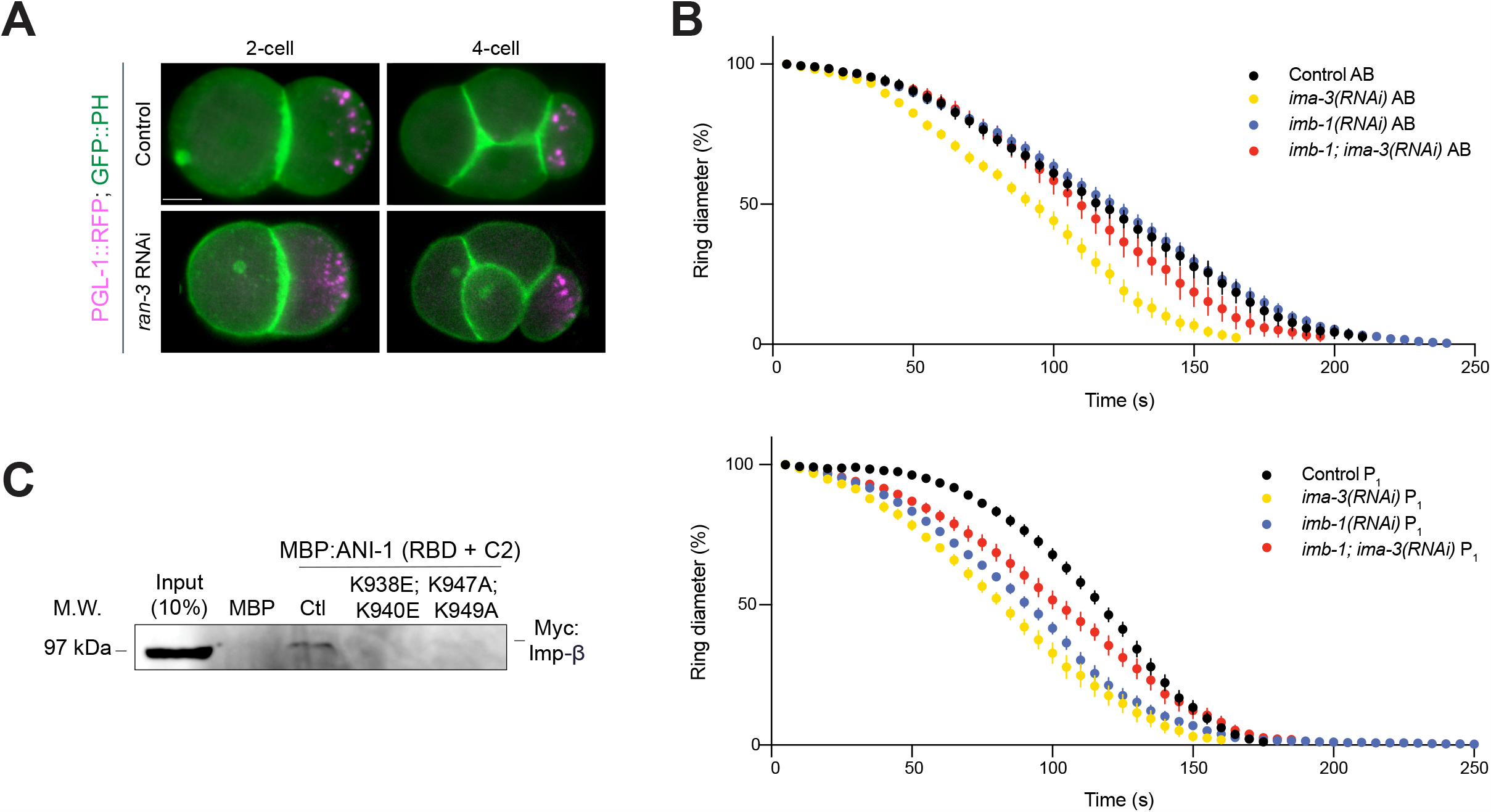
The Ran pathway functions through importins in AB and P_1_ cells. A) Images show embryos co-expressing PGL-1:RFP and GFP::PH in 2-cell and 4-cell control (top) and *ran-3(RNAi)* (bottom) embryos. B) A graph shows ring closure in AB (top) and P_1_(bottom) *ima-3(RNAi)* (yellow), *imb-1(RNAi)* (blue), *imb-1(RNAi); ima-3(RNAi)* (red) and control (AB, coral; P_1_, black) cells. C) A western blot shows Myc-tagged importin-β from HeLa cell lysates (input) and after pull-down with recombinant, purified MBP or MBP-tagged ANI-1 (RBD + C2) containing mutations K938E; K940E or K947A; K949A. All error bars show SEM.

**Movie 1. HILO imaging of the AB cell**. Timelapse images show GFP::NMY-2 at the cortical surface of the AB cell during cytokinesis. Images were recorded every 2 seconds, and the video playback rate is at 7 frames per second. Scale bar is 10 µm.

**Movie 2. HILO imaging of the P**_**1**_ **cell**. Timelapse images show GFP::NMY-2 at the cortical surface of the P_1_ cell during cytokinesis. Images were recorded every 2 seconds, and the video playback rate is at 7 frames per second. Scale bar is 10 µm.

**Movie 3. HILO imaging of AB and P**_**1**_ **cells after *ect-2(RNAi)***. Timelapse images show GFP::NMY-2 at the cortical surface of AB (top) and P_1_ (bottom) cells after *ect-2(RNAi)* during cytokinesis. Images were recorded every 2 seconds, video playback rate is at 7 frames per second. Scale bar is 10 µm.

**Movie 4. Confocal imaging of AB and P**_**1**_ **cells that fail cytokinesis after *ect-2(RNAi)***. Timelapse images show GFP::NMY-2 at the cortical surface of AB (top) and P_1_ (bottom) cells that fail cytokinesis after *ect-2(RNAi)*. Images were recorded every 2 seconds, video playback rate is at 7 frames per second. Scale bar is 10 µm.

**Movie 5. HILO imaging of P**_**0**_ **daughter cells after *par-1(RNAi)***. Timelapse images show GFP::NMY-2 at the cortical surface of a P_0_ daughter cell after *par-1(RNAi)* during cytokinesis. Images were recorded every 2 seconds, video playback rate is at 7 frames per second. Scale bar is 10 µm.

**Movie 6. HILO imaging of tetraploid AB and P**_**1**_ **cells**. Timelapse images show GFP::NMY-2 at the cortical surface of tetraploid AB (top) and P_1_ (bottom) cells during cytokinesis. Images were recorded every 2 seconds, video playback rate is at 7 frames per second. Scale bar is 10 µm.

**Movie 7. HILO imaging of an AB cell after *ran-3(RNAi)***. Timelapse images show GFP::NMY-2 at the cortical surface of an AB cell after *ran-3(RNAi)* during cytokinesis. Images were recorded every 2 seconds, and the video playback rate is at 7 frames per second. Scale bar is 10 µm.

**Movie 8. HILO imaging of a P**_**1**_ **cell after *ran-3(RNAi)***. Timelapse images show GFP::NMY-2 at the cortical surface of a P_1_ cell after *ran-3(RNAi)* during cytokinesis. Images were recorded every 2 seconds, and the video playback rate is at 7 frames per second. Scale bar is 10 µm.

## Uncategorized References

1. Askjaer, P., Galy, V., Hannak, E., & Mattaj, I. W. (2002, Dec). Ran GTPase cycle and importins alpha and beta are essential for spindle formation and nuclear envelope assembly in living Caenorhabditis elegans embryos. Mol Biol Cell, 13(12), 4355–4370. https://doi.org/10.1091/mbc.e02-06-0346

2. Beaudet, D., Akhshi, T., Phillipp, J., Law, C., & Piekny, A. (2017, Nov 15). Active Ran regulates anillin function during cytokinesis. Mol Biol Cell, 28(24), 3517–3531. https://doi.org/10.1091/mbc.E17-04-0253

3. Beaudet, D., Pham, N., Skaik, N., & Piekny, A. (2020, May 15). Importin binding mediates the intramolecular regulation of anillin during cytokinesis. Mol Biol Cell, 31(11), 1124-1139. https://doi.org/10.1091/mbc.E20-01-0006

4. Brenner, S. (1974). The genetics of Caenorhabditis elegans. Genetics, 77(1), 71–94. https://doi.org/10.1002/cbic.200300625

5. Cabernard, C., Prehoda, K. E., & Doe, C. Q. (2010, Sep 2). A spindle-independent cleavage furrow positioning pathway. Nature, 467(7311), 91–94. https://doi.org/10.1038/nature09334

6. Carpentier, G. (2010). Zoom in Images and Stacks. Retrieved August 18 2020 from https://imagej.nih.gov/ij/macros/tools/Zoom_in_Images_and_Stacks.txt

7. Carvalho, A., Desai, A., & Oegema, K. (2009, May 29). Structural memory in the contractile ring makes the duration of cytokinesis independent of cell size. Cell, 137(5), 926–937. https://doi.org/10.1016/j.cell.2009.03.021

8. Chalamalasetty, R. B., Hummer, S., Nigg, E. A., & Sillje, H. H. (2006, Jul 15). Influence of human Ect2 depletion and overexpression on cleavage furrow formation and abscission. J Cell Sci, 119(Pt 14), 3008–3019. https://doi.org/10.1242/jcs.03032

9. Chan, F. Y., Silva, A. M., Saramago, J., Pereira-Sousa, J., Brighton, H. E., Pereira, M., Oegema, K., Gassmann, R., & Carvalho, A. X. (2019, Jan 1). The ARP2/3 complex prevents excessive formin activity during cytokinesis. Mol Biol Cell, 30(1), 96–107. https://doi.org/10.1091/mbc.E18-07-0471

10. Clarke, E. K., Gomez, K. A. R., Mustachi, Z., Murph, M. C., & Schvarzstein, M. (2018). Manipulation of Ploidy in Caenorhabditis elegans. (March), 1-11. https://doi.org/10.3791/57296

11. Clarke, P. R., & Zhang, C. (2008, Jun). Spatial and temporal coordination of mitosis by Ran GTPase. Nat Rev Mol Cell Biol, 9(6), 464–477. https://doi.org/10.1038/nrm2410

12. Cowan, C. R., & Hyman, A. A. (2007, Mar). Acto-myosin reorganization and PAR polarity in C. elegans. Development, 134(6), 1035–1043. https://doi.org/10.1242/dev.000513

13. Dechant, R., & Glotzer, M. (2003). Centrosome Separation and Central Spindle Assembly Act in Redundant Pathways that Regulate Microtubule Density and Trigger Cleavage Furrow Formation. Dev Cell, 4(3), 333–344. https://doi.org/10.1016/s1534-5807(03)00057-1

14. Deng, M., Suraneni, P., Schultz, R. M., & Li, R. (2007, Feb). The Ran GTPase mediates chromatin signaling to control cortical polarity during polar body extrusion in mouse oocytes. Dev Cell, 12(2), 301–308. https://doi.org/10.1016/j.devcel.2006.11.008

15. Echard, A., & O’Farrell, P. H. (2003, Mar 4). The degradation of two mitotic cyclins contributes to the timing of cytokinesis. Curr Biol, 13(5), 373–383. https://doi.org/10.1016/s0960-9822(03)00127-1

16. Evans, T. C. (2006). Transformation and microinjection. WormBook. https://doi.org/doi/10.1895/wormbook.1.108.1

17. Geles, K. G., & Adam, S. A. (2001, May). Germline and developmental roles of the nuclear transport factor importin alpha3 in C. elegans. Development, 128(10), 1817–1830. https://www.ncbi.nlm.nih.gov/pubmed/11311162

18. Green, R. A., Paluch, E., & Oegema, K. (2012). Cytokinesis in animal cells. Annu Rev Cell Dev Biol, 28(1), 29–58. https://doi.org/10.1146/annurev-cellbio-101011-155718

19. Hasegawa, K., Ryu, S. J., & Kalab, P. (2013, Jan 21). Chromosomal gain promotes formation of a steep RanGTP gradient that drives mitosis in aneuploid cells. J Cell Biol, 200(2), 151–161. https://doi.org/10.1083/jcb.201206142

20. Illukkumbura, R., Bland, T., & Goehring, N. W. (2020, Feb). Patterning and polarization of cells by intracellular flows. Curr Opin Cell Biol, 62, 123–134. https://doi.org/10.1016/j.ceb.2019.10.005

21. Kalab, P., Pralle, A., Isacoff, E. Y., Heald, R., & Weis, K. (2006, Mar 30). Analysis of a RanGTP-regulated gradient in mitotic somatic cells. Nature, 440(7084), 697–701. https://doi.org/10.1038/nature04589

22. Kalab, P., Weis, K., & Heald, R. (2002, Mar 29). Visualization of a Ran-GTP gradient in interphase and mitotic Xenopus egg extracts. Science, 295(5564), 2452–2456. https://doi.org/10.1126/science.1068798

23. Kamath, R. S., Martinez-Campos, M., Zipperlen, P., Fraser, A. G., & Ahringer, J. (2001). Effectiveness of specific RNA-mediated interference through ingested double-stranded RNA in Caenorhabditis elegans. Genome biology, 2(1), 1–10.

24. Kemphues, K. J., Priess, J. R., Morton, D. G., & Cheng, N. S. (1988, Feb 12). Identification of genes required for cytoplasmic localization in early C. elegans embryos. Cell, 52(3), 311–320. https://doi.org/10.1016/s0092-8674(88)80024-2

25. Khaliullin, R. N., Green, R. A., Shi, L. Z., Gomez-Cavazos, J. S., Berns, M. W., Desai, A., & Oegema, K. (2018, Jul 2). A positive-feedback-based mechanism for constriction rate acceleration during cytokinesis in Caenorhabditis elegans. Elife, 7. https://doi.org/10.7554/eLife.36073

26. Kiyomitsu, T., & Cheeseman, I. M. (2013, Jul 18). Cortical dynein and asymmetric membrane elongation coordinately position the spindle in anaphase. Cell, 154(2), 391–402. https://doi.org/10.1016/j.cell.2013.06.010

27. Kotynkova, K., Su, K. C., West, S. C., & Petronczki, M. (2016, Dec 6). Plasma Membrane Association but Not Midzone Recruitment of RhoGEF ECT2 Is Essential for Cytokinesis. Cell Rep, 17(10), 2672–2686. https://doi.org/10.1016/j.celrep.2016.11.029

28. Lewellyn, L., Dumont, J., Desai, A., & Oegema, K. (2010, Jan 1). Analyzing the effects of delaying aster separation on furrow formation during cytokinesis in the Caenorhabditis elegans embryo. Mol Biol Cell, 21(1), 50–62. https://doi.org/10.1091/mbc.E09-01-0089

29. Maddox, A. S., Lewellyn, L., Desai, A., & Oegema, K. (2007, May). Anillin and the septins promote asymmetric ingression of the cytokinetic furrow. Dev Cell, 12(5), 827–835. https://doi.org/10.1016/j.devcel.2007.02.018

30. Munro, E., Nance, J., & Priess, J. R. (2004, Sep). Cortical flows powered by asymmetrical contraction transport PAR proteins to establish and maintain anterior-posterior polarity in the early C. elegans embryo. Dev Cell, 7(3), 413–424. https://doi.org/10.1016/j.devcel.2004.08.001

31. Nishimura, Y., & Yonemura, S. (2006, Jan 1). Centralspindlin regulates ECT2 and RhoA accumulation at the equatorial cortex during cytokinesis. J Cell Sci, 119 (Pt 1), 104–114. https://doi.org/10.1242/jcs.02737

32. Osorio, D. S., Chan, F. Y., Saramago, J., Leite, J., Silva, A. M., Sobral, A. F., Gassmann, R., & Carvalho, A. X. (2019, Nov 12). Crosslinking activity of non-muscle myosin II is not sufficient for embryonic cytokinesis in C. elegans. Development, 146 (21). https://doi.org/10.1242/dev.179150

33. Ozugergin, I., & Piekny, A. (2020, Feb 14). Complementary functions for the Ran gradient during division. Small GTPases, 1–11. https://doi.org/10.1080/21541248.2020.1725371

34. Petronczki, M., Glotzer, M., Kraut, N., & Peters, J. M. (2007, May). Polo-like kinase 1 triggers the initiation of cytokinesis in human cells by promoting recruitment of the RhoGEF Ect2 to the central spindle. Dev Cell, 12(5), 713–725. https://doi.org/10.1016/j.devcel.2007.03.013

35. Piekny, A., Werner, M., & Glotzer, M. (2005, Dec). Cytokinesis: welcome to the Rho zone. Trends Cell Biol, 15(12), 651–658. https://doi.org/10.1016/j.tcb.2005.10.006

36. Piekny, A. J., & Glotzer, M. (2008, Jan 8). Anillin is a scaffold protein that links RhoA, actin, and myosin during cytokinesis. Curr Biol, 18(1), 30–36. https://doi.org/10.1016/j.cub.2007.11.068

37. Piekny, A. J., & Maddox, A. S. (2010, Dec). The myriad roles of Anillin during cytokinesis. Semin Cell Dev Biol, 21(9), 881–891. https://doi.org/10.1016/j.semcdb.2010.08.002

38. Pimpale, L. G., Middelkoop, T. C., Mietke, A., & Grill, S. W. (2020, Jul 9). Cell lineage-dependent chiral actomyosin flows drive cellular rearrangements in early Caenorhabditis elegans development. Elife, 9. https://doi.org/10.7554/eLife.54930

39. Pollard, T. D., & O’Shaughnessy, B. (2019, Jun 20). Molecular Mechanism of Cytokinesis. Annu Rev Biochem, 88, 661–689. https://doi.org/10.1146/annurev-biochem-062917-012530

40. Price, K. L., & Rose, L. S. (2017, Sep 1). LET-99 functions in the astral furrowing pathway, where it is required for myosin enrichment in the contractile ring. Mol Biol Cell, 28(18), 2360–2373. https://doi.org/10.1091/mbc.E16-12-0874

41. Prokopenko, S. N., Brumby, A., O’Keefe, L., Prior, L., He, Y., Saint, R., & Bellen, H. J. (1999, Sep 1). A putative exchange factor for Rho1 GTPase is required for initiation of cytokinesis in Drosophila. Genes Dev, 13(17), 2301–2314. https://doi.org/10.1101/gad.13.17.2301

42. Reymann, A. C., Staniscia, F., Erzberger, A., Salbreux, G., & Grill, S. W. (2016, Oct 10). Cortical flow aligns actin filaments to form a furrow. Elife, 5. https://doi.org/10.7554/eLife.17807

43. Rodrigues, N. T., Lekomtsev, S., Jananji, S., Kriston-Vizi, J., Hickson, G. R., & Baum, B. (2015, Aug 27). Kinetochore-localized PP1-Sds22 couples chromosome segregation to polar relaxation. Nature, 524(7566), 489–492. https://doi.org/10.1038/nature14496

44. Rose, L., & Gonczy, P. (2014, Dec 30). Polarity establishment, asymmetric division and segregation of fate determinants in early C. elegans embryos. WormBook, 1-43. https://doi.org/10.1895/wormbook.1.30.2

45. Silva, A. M., Osorio, D. S., Pereira, A. J., Maiato, H., Pinto, I. M., Rubinstein, B., Gassmann, R., Telley, I. A., & Carvalho, A. X. (2016, Dec 19). Robust gap repair in the contractile ring ensures timely completion of cytokinesis. J Cell Biol, 215(6), 789–799. https://doi.org/10.1083/jcb.201605080

46. Silverman-Gavrila, R. V., Hales, K. G., & Wilde, A. (2008, Sep). Anillin-mediated targeting of peanut to pseudocleavage furrows is regulated by the GTPase Ran. Mol Biol Cell, 19(9), 3735–3744. https://doi.org/10.1091/mbc.E08-01-0049

47. Singh, D., & Pohl, C. (2014, Feb 10). Coupling of rotational cortical flow, asymmetric midbody positioning, and spindle rotation mediates dorsoventral axis formation in C. elegans. Dev Cell, 28(3), 253–267. https://doi.org/10.1016/j.devcel.2014.01.002

48. Strome, S., & Wood, W. B. (1982, Mar). Immunofluorescence visualization of germ-line-specific cytoplasmic granules in embryos, larvae, and adults of Caenorhabditis elegans. Proc Natl Acad Sci U S A, 79(5), 1558–1562. https://doi.org/10.1073/pnas.79.5.1558

49. Tatsumoto, T., Xie, X., Blumenthal, R., Okamoto, I., & Miki, T. (1999, Nov 29). Human ECT2 is an exchange factor for Rho GTPases, phosphorylated in G2/M phases, and involved in cytokinesis. J Cell Biol, 147(5), 921–928. https://doi.org/10.1083/jcb.147.5.921

50. Tokunaga, M., Imamoto, N., & Sakata-Sogawa, K. (2008, Feb). Highly inclined thin illumination enables clear single-molecule imaging in cells. Nat Methods, 5(2), 159–161. https://doi.org/10.1038/nmeth1171

51. Tse, Y. C., Piekny, A., & Glotzer, M. (2011, Sep). Anillin promotes astral microtubule-directed cortical myosin polarization. Mol Biol Cell, 22(17), 3165–3175. https://doi.org/10.1091/mbc.E11-05-0399

52. Tse, Y. C., Werner, M., Longhini, K. M., Labbe, J. C., Goldstein, B., & Glotzer, M. (2012, Oct). RhoA activation during polarization and cytokinesis of the early Caenorhabditis elegans embryo is differentially dependent on NOP-1 and CYK-4. Mol Biol Cell, 23(20), 4020–4031. https://doi.org/10.1091/mbc.E12-04-0268

53. van Oostende Triplet, C., Jaramillo Garcia, M., Haji Bik, H., Beaudet, D., & Piekny, A. (2014, Sep 1). Anillin interacts with microtubules and is part of the astral pathway that defines cortical domains. J Cell Sci, 127(Pt 17), 3699–3710. https://doi.org/10.1242/jcs.147504

54. Xu, L., & Massague, J. (2004, Mar). Nucleocytoplasmic shuttling of signal transducers. Nat Rev Mol Cell Biol, 5(3), 209–219. https://doi.org/10.1038/nrm1331

55. Yuce, O., Piekny, A., & Glotzer, M. (2005, Aug 15). An ECT2-centralspindlin complex regulates the localization and function of RhoA. J Cell Biol, 170(4), 571–582. https://doi.org/10.1083/jcb.200501097

56. Zanin, E., Desai, A., Poser, I., Toyoda, Y., Andree, C., Moebius, C., Bickle, M., Conradt, B., Piekny, A., & Oegema, K. (2013, Sep 16). A conserved RhoGAP limits M phase contractility and coordinates with microtubule asters to confine RhoA during cytokinesis. Dev Cell, 26(5), 496–510. https://doi.org/10.1016/j.devcel.2013.08.005

57. Zhao, W. M., & Fang, G. (2005, Sep 13). MgcRacGAP controls the assembly of the contractile ring and the initiation of cytokinesis. Proc Natl Acad Sci U S A, 102(37), 13158–13163. https://doi.org/10.1073/pnas.0504145102

